# Dairy practices select for thermostable endolysins in phages

**DOI:** 10.1101/2023.09.27.559745

**Authors:** Frank Oechslin, Xiaojun Zhu, Carlee Morency, Vincent Somerville, Rong Shi, Sylvain Moineau

## Abstract

Endolysins are phage lytic enzymes that degrade the bacterial cell wall to release phage progeny. We found that the diversity of endolysins is unsuspectedly low and dominated by a single structural type in distinct phages that infect *Streptococcus thermophilus*. Through X-ray crystallography, we discovered that this type of endolysin contains a highly conserved calcium-binding motif in its cell wall binding domain. Inactivation of the motif in the purified endolysin or in the phage genome revealed its key role in stabilizing the enzyme under conditions that mimic the cheesemaking process, including at elevated temperatures. Fermented dairy products such as yogurt and cheeses are incubated or cooked at higher temperatures and require the addition of thermophilic lactic acid bacteria such as *S. thermophilus* to drive milk fermentation. It appears that dairy practices have influenced the genetic signatures of streptococcal phages by selecting for thermostable endolysins, which in turn have limited their diversity.

## Introduction

Bacteriophages, also referred to as phages, are bacterial viruses that are found in the same ecological niches as their hosts. They are highly diverse and are now classified into numerous viral families based on their genome composition ^1,2^. Phages play a pivotal role in regulating bacterial populations and shaping their evolution through selective pressure and gene transfer ^3^. They can also serve as biocontrol agents to prevent or suppress bacterial infections in plants, animals, and humans and are being actively considered as an alternative or complement to antibiotics in response to multi-drug- resistant infections ^4^. However, for decades, phages have also been considered “enemies” in various industrial processes where they sometimes kill bacteria that play a beneficial role in fermentation or in biotechnology. For instance, phage-induced lysis of starter cultures remains the primary cause of milk fermentation failure in the cheese industry, resulting in low-quality products ^5^.

The ability of phages to lyse their bacterial host at the end of their lytic cycle involves the coordinated action of at least two types of viral proteins: holins and endolysins^6^. The holin protein is synthesized intracellularly to generate pores in the bacterial plasma membrane. These pores allow the endolysin, a peptidoglycan hydrolase, to enter the extracytoplasmic space and degrade the cell wall, leading to bacterial lysis and the release of new virions ^7^. In some cases, the endolysins may function independently of holins and rely on the host secretory pathway to migrate across the plasma membrane ^8^.

Phage endolysins target the bacterial peptidoglycan, a vital structure composed of a complex meshwork of N-acetylglucosamine (GlcNAc)-N-acetylmuramic acid (MurNAc) glycan strands that are cross-linked by short stem peptides attached to MurNac residues^9^. While all phage endolysins have the same function of cleaving peptidoglycan, their structure and mode of action can differ significantly. In general, endolysins that target Gram-positive bacteria have a modular structure that includes one or more enzymatically active domain (EAD) connected to a cell wall-binding domain (CBD) by a short linker ^10^. Based on the peptidoglycan bond they hydrolyze, endolysins are currently classified into four primary groups. These include glucosaminidases and muramidases that hydrolyze glycan chains as well as amidases and endopeptidases that hydrolyze the peptidoglycan stem peptides or interconnecting bridges. The CBD domain provides high bacterial specificity and helps optimally position the enzyme to degrade its peptidoglycan substrate ^11,12^. Overall, EAD and CBD can be very different, and their various combinations have resulted in a high degree of diversity throughout phage evolution ^13,14^.

The reason for the diversity of endolysins is not yet fully understood, but is likely related to the adaptation of phages to diverse and changing bacterial cell wall compositions ^15^. Additionally, endolysin domains may have evolved to target essential cell wall factors for host viability ^16^ or as an adaptation to preferred or new phage hosts ^11^. In a recent study, we investigated the diversity of endolysins found in phages infecting the Gram-positive dairy bacterium *Lactococcus lactis* ^17^. Our results showed significant diversity, revealing 11 structural types associated with specific phage genera. Furthermore, we demonstrated that this endolysin diversity is a result of host adaptation, rather than being dependent on the type of phage genera. Indeed, endolysin genes could be exchanged between phages from different genera with minimal fitness costs, as long as both the donor and recipient phages infected the same bacterial strain.

In this study, we aimed to investigate the diversity of endolysins present in phages infecting *Streptococcus thermophilus*, which is considered the second most important dairy bacterial species after *L. lactis* ^18^. We found that the diversity of endolysins is significantly lower than in lactococcal phages and is not restricted to specific phage genera. In fact, over 90% of the endolysins belonged to only one structural type and it is found in phages belonging to 4 of the 5 known phage genera infecting *S. thermophilus*. Structural analysis revealed that this type of endolysin possesses an N-terminal catalytic amidase_5 domain and a highly conserved calcium binding motif (CaBM) in the CBD. Targeted mutagenesis of CaBM in the purified endolysin or its gene in the phage genome revealed its importance in stabilizing the enzyme at elevated temperatures. Indeed, CaBM increased the thermal stability of the endolysin and its inactivation in the phage genome resulted in temperature-dependent fitness costs and phages could not resume their lytic cycle upon heat shock stress. Several cheeses are produced at elevated temperatures and require thermophilic lactic acid bacteria such as *S. thermophilus* ^19^. Humans have domesticated *S. thermophilus* for centuries if not more, a process that has left genetic hallmarks such as the loss of specific bacterial genes ^20^. It now appears that practices such as cheese production have also influenced the genetic signatures of streptococcal phages by selecting for thermostable endolysins and limiting their diversity.

## Results

### Endolysins of phages that infect *S. thermophilus* can be classified into five types

To assess the diversity of endolysins present in streptococcal phages, we first retrieved all 195 phage genomes infecting *S. thermophilus* from NCBI and analyzed their phylogenetic relatedness and gene conservation (Figure 1). The phage genomes represented the five phage genera infecting *S. thermophilus*, namely, *Moineauvirus* (former *cos*), *Brussowvirus* (former *pac*), *Vansinderenvirus (*former 5093), 987, and P738. Based on HHPred, BLASTP, and AlphaFold, we predicted five types of endolysin structures. Group A endolysins, with an average predicted molecular weight of 30.8 ± 0.3 kDa, were by far the most abundant as they were found in 91% (n = 178) of the phage genomes. They all have a predicted N-terminal catalytic amidase_5 domain that belong to the N1pC/P60 family of cell wall peptidases and a C-terminal region homologous to the target recognition domain of zoocin A, a Zn-metallopeptidase secreted by *Streptococcus zooepidemicus* ^21,22^. The EAD domain of group A endolysin also belongs to the cysteine/histidine-dependent amidohydrolase/peptidase family (CHAP), as the characteristic cysteine (C35) and histidine (H101) residues were conserved in all of them (Supplementary Figure 1) ^23^. These residues were at a nearly similar position in the amidase_5 domain observed in the pal endolysin (C34 and H99), an N-acetylmuramoyl-L-alanine amidase encoded by the pneumococcal phage Dp-1 ^24^. Furthermore, we found that half of the type A endolysin genes were disrupted by a group IA2 intron, as reported previously ^25^.

**Figure 1.**
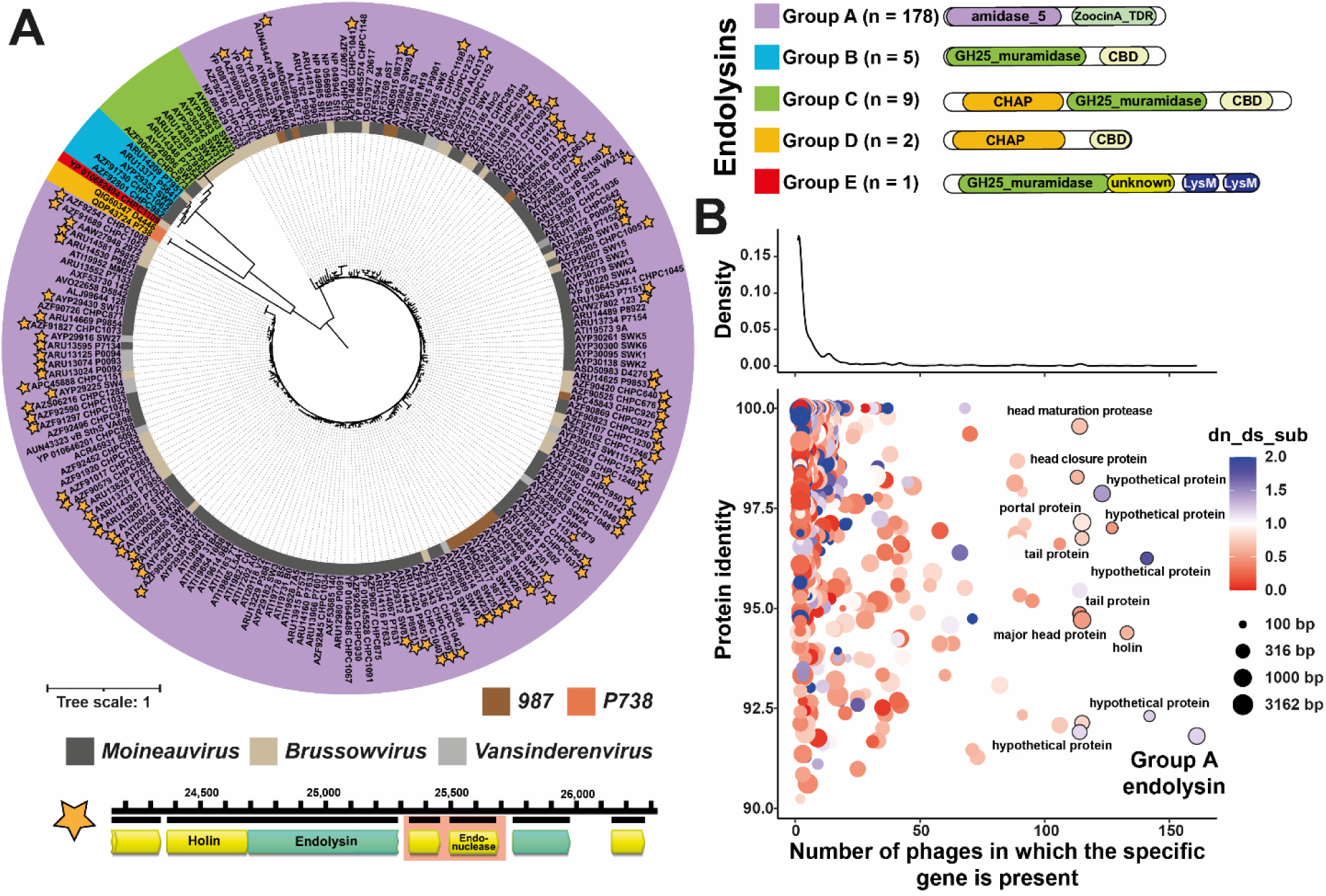
Phylogenetic relationship of endolysins from phages that infect *S. thermophilus*. A set of 195 complete phage genomes infecting *S. thermophilus*, available on NCBI as of February 26, 2023, were analyzed for endolysin diversity. Multiple sequence alignments were performed using MUSCLE (v3.8.425), and the phylogenetic tree was generated using Neighbor-Joining and a Jukes-Cantor genetic distance model. Conserved enzymatically active domain (EAD) and cell wall binding domains (CBD) were identified using HHPred, BLASTP, and AlphaFold predictions. Endolysins were classified into five groups based on their catalytic domains and are represented by different colors. Accession numbers for each endolysin are followed by the name of the phage from which it was found, and each phage genus is depicted by a unique color. The presence of a group IA2 intron within the endolysin gene is indicated with a star, and the intron found in the endolysin gene of phage 2972 (*Brussowvirus*) is provided as an example. B) The gene conservation of all genes in the *S. thermophilus* phage genomes is depicted through protein identity and the number of phages that contain each gene (density plot on the x-axis). Additionally, the dn/ds substitution rate and gene size are represented by the point color and size, respectively. The ten most prevalent genes are annotated for reference.

Group B endolysins (2.6%, n = 5, 32.8 ± 1.0 kDa) were predicted to be muramidases containing a glycosyl hydrolase family 25 (GH25) catalytic domain and a C-terminal region that lacked homology with known CBDs. Endolysins from group C (4.6%, n = 9, 50.8 ± 0.1 kDa) had the same predicted overall architecture as those in group B but with an additional N-terminal CHAP domain that showed homology with the *Enterococcus* phage endolysin LysIME-EF1 ^26^. Group D endolysins (1.0%, n = 2, 27.4 ± 0.1 kDa) were rare but also had an N-terminal CHAP domain homologous to LysIME-EF1 endolysins but with a C-terminal domain without homology with known CBDs. Finally, the group E endolysin 0.5%, n = 1, 45.55 kDa) contained only one member, namely the one from the phage CHPC1109. This very rare endolysin was predicted to have a GH25 EAD muramidase and two additional C-terminal LysM domains. AlphaFold structure prediction revealed an additional third domain of unknown function that was not predicted by either Blast or HHpred (Supplementary Figure 2).

We observed no clear correlation between the phylogeny of endolysins and the taxonomy of streptococcal phages, except for group D endolysins, which were exclusively associated to P738-like phages. Type B and C endolysins were shared between *Brussowvirus* and *Moineauvirus*, while type A endolysins were distributed across all phage genera except P738. Notably, type A endolysins were the most prevalent (n=178) among all *S. thermophilus* phage genomes even when compared to other conserved non-structural proteins like holin (n=133), or structural proteins such as major head (n=115), tail (n=114), or portal (n=115) proteins. However, despite the numerical dominance of type A endolysins, the other genes (holin, major head, tail, and portal proteins) appeared to be more functionally constrained, as indicated by their lower ratio of non-synonymous to synonymous substitutions (dN/dS) (Figure 1B).

### Structural analysis of Lys2972 revealed that it possesses a unique metal-binding motif that is highly conserved among group A endolysins

To gain a more comprehensive understanding of the characteristics contributing to the prevalence of type A endolysins among streptococcal phages, a structural analysis of the endolysin from phage 2972 (Lys2972) was performed. This endolysin was selected as a representative of type A, and its crystal structure was determined in the C2 space group, with one molecule per asymmetric unit. All residues, except for 235-237 and the final three residues (278-280) at the C-terminal position, were successfully built.

Lys2972 (280 amino acids) was found to have two main domains, namely an EAD (residues 1-149) and a CBD (155-280), both of which have a globular shape (Figure 2A). In addition, Lys2972 had a relatively short and rigid linker segment (residues 150-154), leading to significant contacts between the two domains and an overall compact structure. A search conducted using the Dali Server revealed a significant similarity between the EAD domain of Lys2972 and other members of the cell wall peptidase superfamily NlpC/p60. Notably, the bacterial peptidoglycan hydrolase SagA from *Enterococcus faecium* (PDB 6B8C) exhibited the highest degree of similarity, with a Z-score of 13.9 and an r.m.s.d. of 2.1 Å for 106 aligned Cα atoms (Supplementary Figure 3A) ^27^. The CBD of Lys2972 was found to have only one structural homolog, corresponding to the target recognition domain (TRD) of bacterial Zoocin A ^22^. The structure of this homolog was determined by NMR (PDB 2LS0) and exhibited a Z-score of 9.8, an rmsd of 2.1 Å for 94 aligned Cα atoms, and a sequence identity of 25.5% (Supplementary Figure 3B).

**Figure 2.**
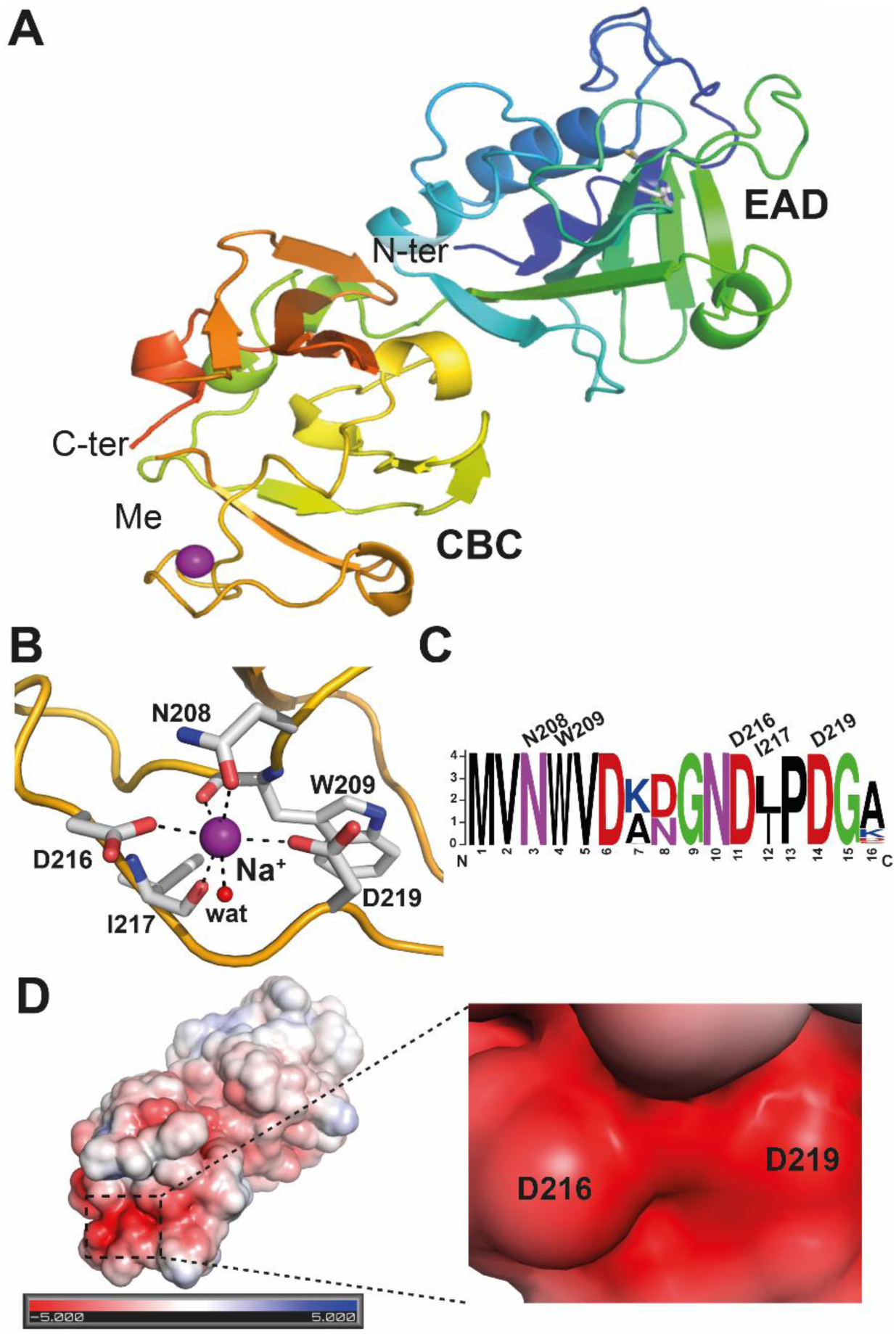
Crystal structure of Lys2972. A) The endolysin is color coded in a rainbow scheme with the N-terminal in blue and the C-terminal in red. Both the EAD and CBD domains are labeled and the metal ion (Na+ or Ca2+) bound to the CBD domain is shown as a purple sphere. B) The interactions between the metal (represented as Na+ in this model) and the main chain carbonyls of W209 and I217, the side chain oxygen atoms of N208, D216, and D219, and a water molecule are depicted as dashed lines. C) Graphical representation of the multiple sequence alignment of the CaBM region found in the 178 type A endoylsin sequences used in this study. D) Surface representation of Lys2972 colored according to its electrostatic potential (ranging from -5 kBT/e (red) to 5 kBT/e (blue)) and calculated using PyMol (Schrödinger). The metal binding site is largely negatively-charged and the enzyme is shown in the same orientation as in panel A.

Interestingly, a metal ion was identified at the edge of the CBD domain and was observed to be coordinated by the carboxylate groups of Asp216 and Asp219, the main chain carbonyl groups of Trp209 and Ile217, as well as the side chain carbonyl of Asn208 and a water molecule (Figure 2B). While this novel metal binding site and its corresponding residues were absent in the previously mentioned Zoocin A TRD structure, they were remarkably conserved in all type A endolysins (Figure 2C).

An electrostatic surface charge calculation further revealed a heavily negatively charged patch at this position (Figure 2D), consistent with its metal binding capacity. Considering that the coordination number of the metal is six, all the atoms in the coordination sphere of this metal are oxygens (coming from carboxylate, carbonyl, or water molecules), and the metal-ligand distances range between 2.2 and 2.5 Å, with an average value of 2.41 Å, this metal is expected to be either calcium or sodium ^28^. Although refinement with Na+ has resulted in a temperature factor more in line with that of the coordinating oxygen atoms, this may be attributed to the low pH (5.0), which diminishes the binding capacity of D216 and D219 for divalent calcium ions, and the high abundance of sodium ions in the crystallization condition (>100 mM). However, it can be presumed that under neutral pH conditions and/or in the presence of high calcium concentration in milk, calcium is likely the preferred ion bound to this metal-binding motif. Consequently, this specific motif was further designated as a Calcium-Binding Motif (CaBM).

### Calcium plays a role in the lytic activity of Lys2972

Following the structural analysis described above, the role of calcium in the lytic activity of Lys2972 was investigated by inactivating the CaBM. The amino acids providing their side chains for metal coordination were replaced by alanine to construct single mutants (Lys2972>N208A, Lys2972>D216A, and Lys2972>D219A), as well as a triple-mutant-designated Lys2972^CaBM^ (Lys2972>N208A+D216A+D219A). After Ni affinity and size exclusion chromatography, the purity and molecular weight of each endolysin was confirmed on 4–12% BisTris gels (Supplementary Figure 4).

The enzymatic activity of each purified enzymes (wild-type and four mutants) was assessed by measuring the reduction in turbidity of a *S. thermophilus* cell suspension in the exponential phase after a 15-minute incubation period with or without CaCl_2_. In the presence of 10 mM CaCl_2_, the lytic activity of Lys2972 increased by 24.4 ± 2.0% (at a protein concentration of 1 µM, Figure 3A). In contrast, the Lys2972^CaBM^ mutant was 32.3 ± 2.4% less active than the wild-type. Of note, an increase in lytic activity was also observed for Lys2972>N208A (11.4 ± 1.3%) and Lys2972>D216A (18.0 ± 2.4%) mutants, but to a lesser extent compared to WT Lys2972 (29.6 ± 1.9%) and at a protein concentration of 0.5 µM (Figure 3B). It is noteworthy that the Lys2972>D219A mutant did not show any increase in enzymatic activity in presence of calcium, indicating that D219 plays a crucial role in calcium binding. Additionally, we observed that the calcium response of Lys2972 reached saturation at 0.5 mM CaCl2, while this was not the case for the Lys2972>N208A and Lys2972>D216A mutants, as their activity gradually increased with calcium concentration. Finally, the addition of EDTA (10 mM) resulted in an 11.5 ± 1.5% decrease in Lys2972 activity (Supplementary Figure 5A).

**Figure 3.**
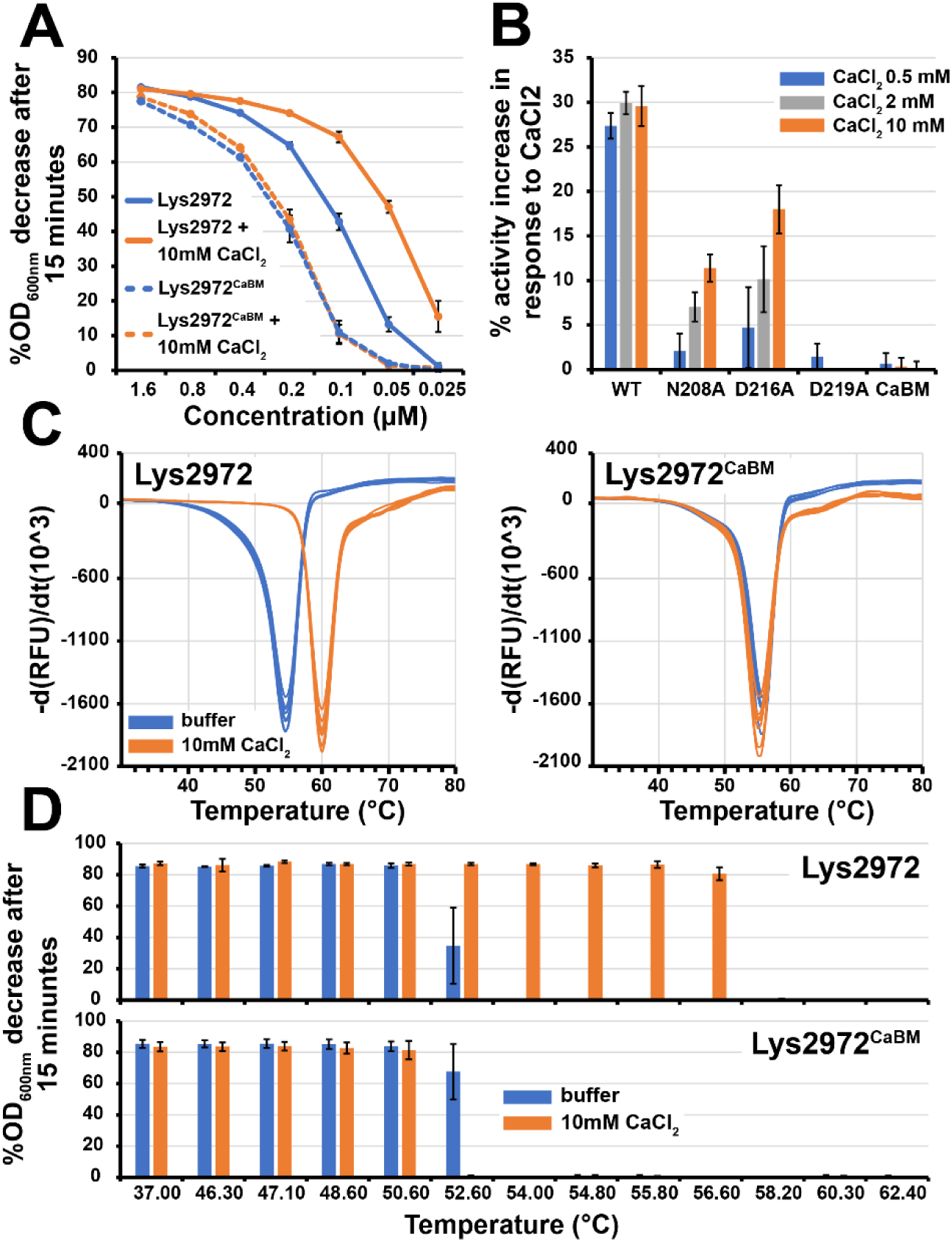
Importance of CaBM for the lytic activity and heat stability of Lys2972. (A) Characterization of the lytic activity of Lys2972 and its inactivated CaBM mutant (Lys2972^CaBM^) at various concentrations and with or without the addition of 10 mM CaCl_2_. The lytic activity was determined by measuring the decrease in turbidity after 15 minutes of *S. thermophilus* DGCC7710 cells in exponential growth phase. **(B)** Characterization of the lytic activity of Lys2972 and CaBM derivative mutants (0.5 uM) after the addition of 0.5, 2, or 10 mM CaCl_2_ compared to values obtained without the addition of CaCl_2_. **(C)** Graph of the melting temperature (Tm) of Lys2972 and Lys2972^CaBM^ with or without the addition of 10 mM CaCl_2_. The thermal shift assay was performed using SYPRO® Orange, and the Tm (lowest part of the curve) was determined by plotting the first derivative of the fluorescence emission as a function of 0.5 °C temperature steps. **(D)** Graph of the thermal stability of Lys2972 and Lys2972^CaBM^ after a 15-minute incubation step at various temperatures and with or without 10 mM CaCl_2_. The lytic activity was determined by measuring the decrease in turbidity after 15 minutes of *S. thermophilus* DGCC7710 cells in the exponential growth phase. Experiments were performed in triplicate.

We also generated a mutant that lacked the entire CBD (Lys2972ΔCBD). This additional mutant exhibited one-hundred-fold reduced activity. Indeed, while Lys2972 achieved approximatively 60% turbidity reduction after 15 minutes at a concentration of 0.2 µM (Figure 3A), the mutant required a concentration of 20 µM to reach comparable activity level (Supplementary Figure 5B). In comparison, the decrease in activity (approx.. 30%) at the same concentration when CaBM was inactivated suggests that its contribution to enzyme activity may not be essential. Instead, CaBM might play a more significant role in other aspects of enzyme function. Other studies have shown that calcium can increase the heat stability of various proteases ^29,30^. Calcium-binding motifs can also play a crucial role in improving mechanical stability by creating additional non-covalent bonds that strengthen the overall protein structure ^29,30^.

### The CaBM is important for the thermal stability of Lys2972

To investigate whether CaBM has a stabilizing effect, we performed a thermal shift assay to compare the stability of Lys2972 and its CaBM mutants. Our results indicated that in the presence of 10 mM CaCl_2_, the melting temperature of Lys2972 increased from 54.5 to 60.0 °C (Figure 3C). However, no such increase was observed with the mutants Lys2972^CaBM^, Lys2972>N208A, Lys2972>D216A, and Lys2972>D219A or when 10 mM of EDTA was added to Lys2972 (Figure 3C and Supplementary Figure 6). The importance of CaBM for the stability of Lys2972 was further substantiated by testing the activity of the endolysin at different temperatures. In the presence of 10 mM CaCl_2_, Lys2972 retained its lytic activity at temperatures up to 56.6 °C, which could not be achieved without CaCl_2_ or an active CaBM (Figure 3D), validating this unique metal-binding motif.

### Inactivation of the CaBM in Lys2972 is associated with a temperature-dependent fitness cost of phage 2972

Since we observed that calcium can increase the thermal stability of Lys2972, we investigated whether temperature could affect the replication of a phage mutant with an inactivated CaBM. This was achieved by replacing the three critical amino acids (N208A, D216A, D219A) responsible for coordinating calcium with alanines using CRISPR-Cas9 genome editing. The resulting phage mutant, designated 2972^CaBM^, produced clear plaques at 37 °C on *S. thermophilus* DGCC7710 (Supplementary Figure 7). Then, we infected the same bacterial host with either phage 2972 or 2972^CaBM^, incubated them at different temperatures, and then measured end turbidity values (Figure 4A). We did not observe any significant differences between 2972 or 2972^CaBM^ under these experimental conditions, as the replication of both phages and the host was impaired at 46 °C and no lytic activity was observed at 49° C. A closer look at the kinetics of cell lysis for phage 2972 or 2972^CaBM^ showed that inhibition of cell lysis occurred for both phages at 47 °C, whereas no differences were observed at 42 °C and 46 °C (Supplementary Figure 8). However, while the lytic plaques produced by phage 2972 (0.34 ± 0.12 mm^2^) and 2972^CaBM^ (0.33 ± 0.11 mm^2^) showed no statistical differences at 37 °C (p-value = 0.1716), the 2972^CaBM^ mutant produced smaller plaques at 42 °C (0.28 ± 0.09 vs. 0.25 ± 0.09 mm^2^, P ≤ 0.001) and at 46 °C (0.17 ± 0.05 vs. 0.11 ± 0.04 mm^2^, P ≤ 0.0001) (Figure 4B). Furthermore, competition assays between 2972 and 2972^CaBM^ revealed a fitness cost associated with the lost of CaBM when the temperature was increased (-0.31 ± 0.03 at 37 °C, -0.57 ± 0.07 at 42 °C, and -0.89 ± 0.08 at 46 °C) (Figure 4C). A linear correlation was also found between relative fitness and plaque size variations between 2972 and 2972^CaBM^ at different temperatures (Figure 4D).

**Figure 4.**
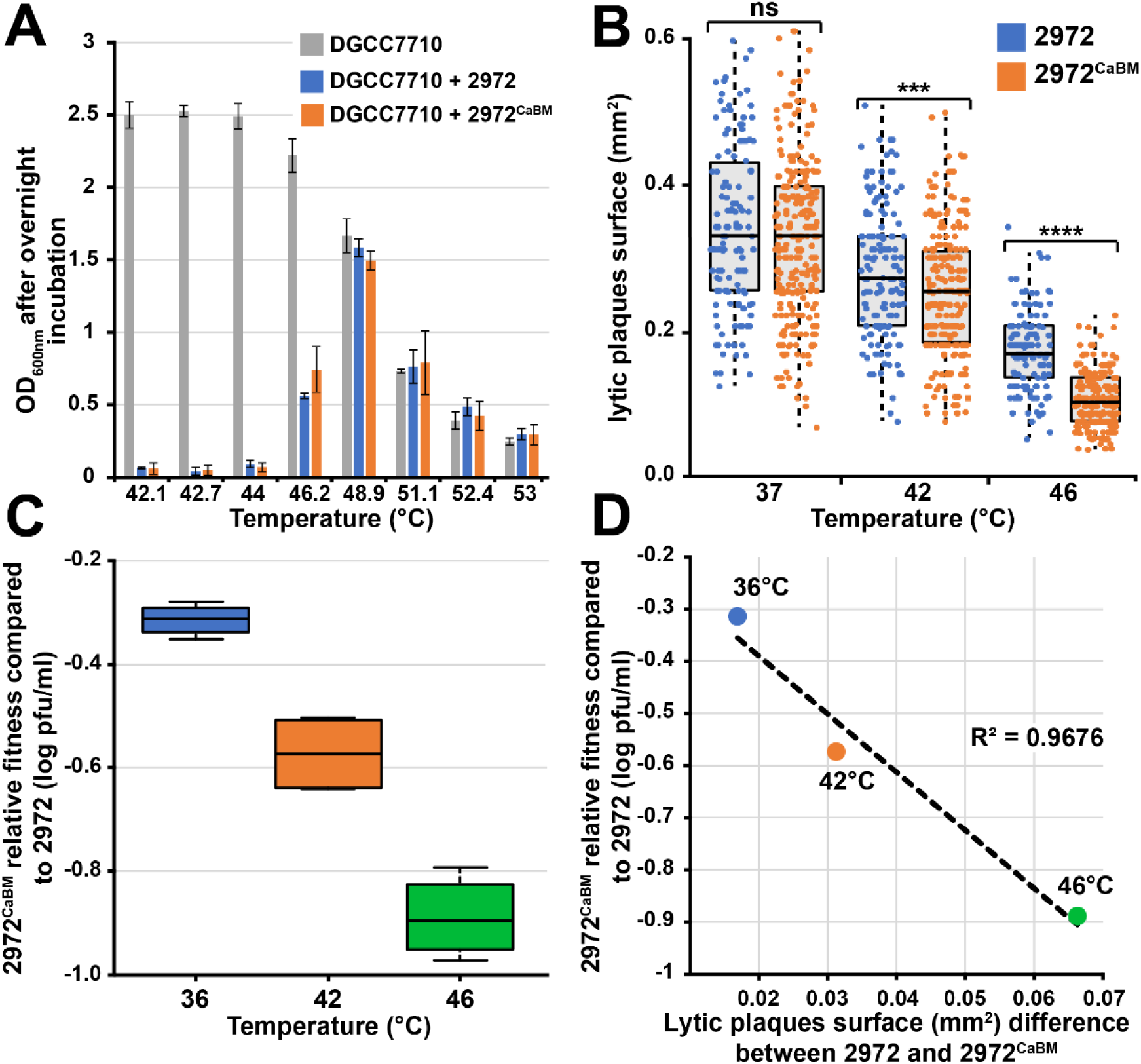
Significance of CaBM in the lytic activity of phage 2972 at different temperatures. (**A)** Lytic activity of phage 2972 and the inactivated CaBM mutant (2972^CaBM^) at different temperatures. *S. thermophilus* DGCC7710 was infected with phage 2972 or 2972^CaBM^ at a MOI of 0.1. Cell culture turbidity was measured after 16 hours of incubation at different temperatures. **(B)** Size of lytic plaques produced by phage 2972 or 2972^CaBM^ when grown at different temperatures (ns = P > 0.05, *** = P ≤ 0.001, **** = P ≤ 0.0001) using Welchʻs two sample t-test). *S. thermophilus* DGCC7710 was used for plaque visualization using a double layer agar assay. **(C)** Impact of CaBM inactivation on phage 2972 fitness. A direct competition assay was performed by co-amplifying both phage 2972 and 2972^CaBM^ at an initial MOI of 0.0001 and at different temperatures. After overnight incubation, the population of phage 2972^CaBM^ was quantified using the strain *S. thermophilus* DGCC7710 BIM/Lys2972 (resistant to phage 2972 but not 2972^CaBM^) and compared with the titer of the total phage population obtained using strain *S. thermophilus* DGCC7710. **(D)** Linear correlation between relative fitness and difference in plaque size between phages 2972 and 2972^CaBM^.

### CaBM enables phage 2972 to resume cell lysis after a heat shock or at a temperature that does not support the growth of its bacterial host

To investigate the role of CaBM on the thermal stability of Lys2972 during phage replication, we also exposed cells infected with either phage 2972 or 2972^CaBM^ to a 15-minute heat shock (60 °C) at different time intervals after infection. A previous study on the global gene expression of phage 2972 revealed that Lys2972 was maximally expressed 27 minutes after infection ^31^. We observed that when infected cells were subjected to heat shock within the first 20 minutes of infection, no significant differences were observed between phages 2972 and 2972^CaBM^ (Figure 5A–5C). In both cases, bacterial lysis was delayed until host growth resumed. However, when the heat shock was applied 30 minutes after infection or later, bacterial lysis occurred only in cells infected with 2972 and not 2972^CaBM^ (Figure 5D–5F). Exposure of infected cells to a 55 °C also revealed a similar trend, albeit with a significantly shorter delay in lysis time for 2972^CaBM^ compared to the 60 °C stress (Supplementary Figure 9).

**Figure 5.**
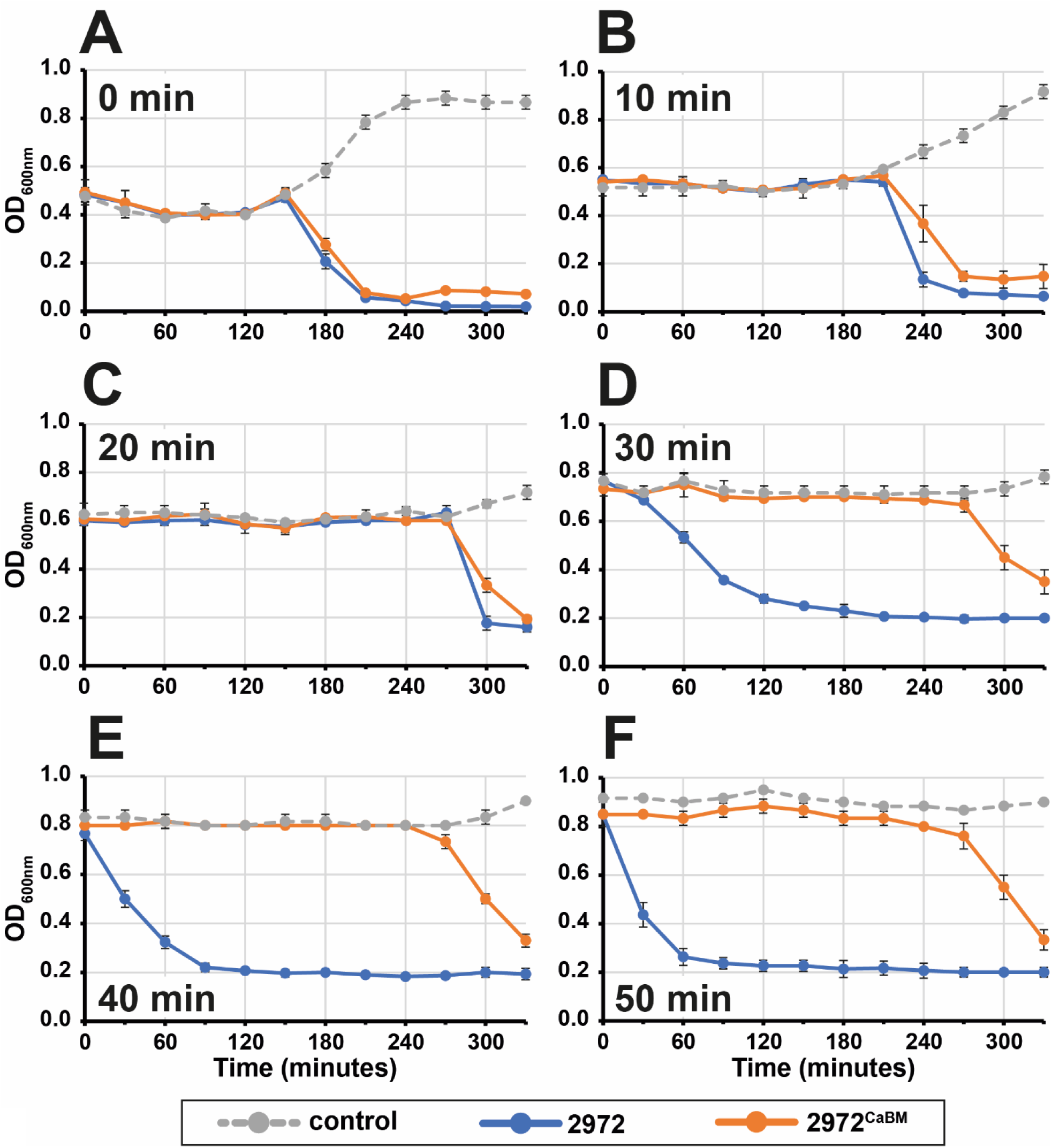
Impact of a 60 °C heat shock on phage 2972 and 2972^CaBM^ lytic activity. *S. thermophilus* DGCC7710 cells in the early exponential phase were infected with phage 2972 or 2972^CaBM^ at a MOI of 0.1. The infected cells were then exposed to a 15-minute heat shock at 60 °C either **(A)** immediately after infection, or **(B-F)** at various time points post-infection. Optical density was measured every 30 minutes after heat shock. Experiments were performed in triplicate.

We also exposed the phage-infected cells to a 150-min incubation period at 55 °C or 60 °C, starting 45 min after infection (Figure 6A–C). The results showed that while a temperature of 55 °C inhibited bacterial growth, phage 2972 was still able to lyse its bacterial host, whereas phage 2972^CaBM^ was not. However, both phages failed to exhibit lytic activity at 60 °C. These results were consistent with the observation that Lys2972 was capable of lysing cells at 55 °C, provided that the CaBM was functional and that calcium was present (as shown in Figure 6D). Overall, the results suggest that CaBM plays a critical role in enhancing its lytic activity at high temperatures during phage 2972 replication. Therefore, CaBM is a key factor in making the lytic cycle of phage 2972 more resilient to temperature fluctuations found in some cheesemaking processes.

**Figure 6.**
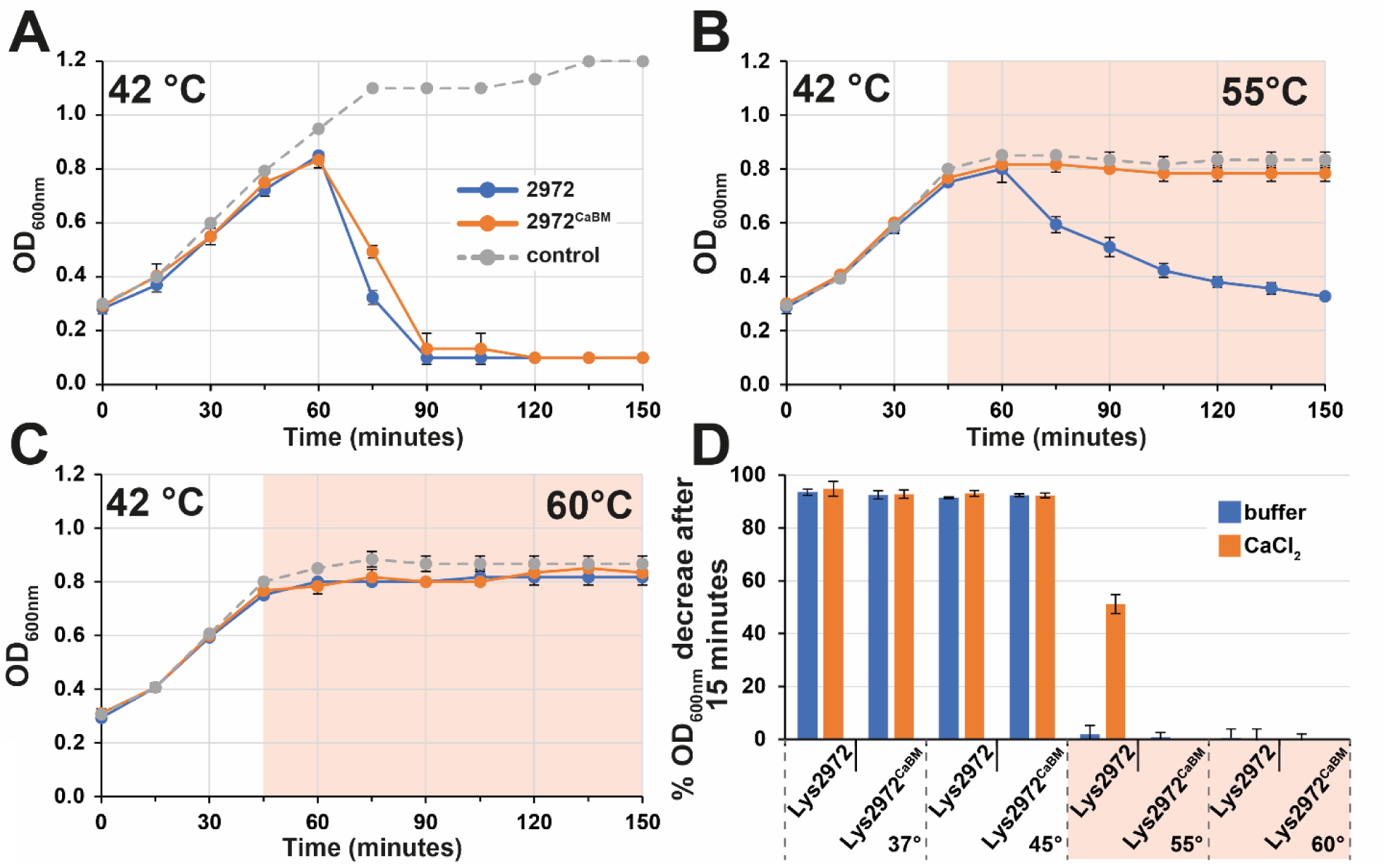
Effect of incubation at 55 °C or 60 °C on the lytic activity of phages 2972 and 2972^CaBM^, as well as endolysins Lys2972 and Lys2972^CaBM^. Cells were infected with phage 2972 or 2972^CaBM^ at a MOI of 0.1 and incubated at either **(A)** 42 °C or **(B)** 42 °C for 45 minutes before switching to 55 °C or **(C)** 60 °C. Optical density was measured every 15 minutes. **(D)** In addition, the activity of the purified endolysin Lys2972 and its derived CaBM mutant (4 µM) was characterized at different temperatures and with or without the addition of 10 mM CaCl^2^. Experiments were performed in triplicate.

### CaBM provides a fitness advantage under thermal conditions used in cheese production

*S. thermophilus* is used as a starter culture in the production of Italian and Swiss hard cheeses ^32^, in which the curd is cooked in whey at temperatures ranging from 45 to 60 °C ^33^. Thus, the fitness of phage 2972^CaBM^ was investigated under the conditions mimicking hard cheese production, i.e.: A) infection of the starter culture at a low MOI ^34^ and B) during temperature fluctuations. Cells were infected at a MOI of 0.0001 and incubated in a programmable water bath with a temperature profile similar to that used in the production of Emmental cheese (Figure 6A) ^35^. This included an initial incubation step that corresponds to maturation of the starter culture, followed by milk coagulation and agitation of the curd. Periods of 60 to 80 minutes were used for the incubation step at 35 °C ^35^. Then, the temperature was raised to 55 °C over 20 minutes, maintained for another 30 minutes, and gradually lowered to 40 °C over 60 minutes to simulate the cooling and pressing of the cheese curd.

Under these thermal conditions, we observed that cell cultures infected with phage 2972 exhibited earlier lysis than those infected with 2972^CaBM^. This was supported by the consistently shorter time at maximum rate (t at max V) of cell lysis kinetics for 2972 compared to 2972^CaBM^ (Figure 7B–D). Conversely, no significant differences were observed between the two phages in the control condition where cells were infected at the same MOI and at a constant temperature of 40 °C (Figure 7E). Finally, the CaBM was again found to provide a selective advantage under these thermal conditions during a competition test between phages 2972 and 2972^CaBM^. The fitness cost associated with the absence of a functional CaBM gradually increased with the duration of the first incubation step at 35 °C (-0.52 ± 0.06 for 60 min, -0.62 ± 0.05 for 70 min, and -0.83 ± 0.1 for 80 min).

**Figure 7.**
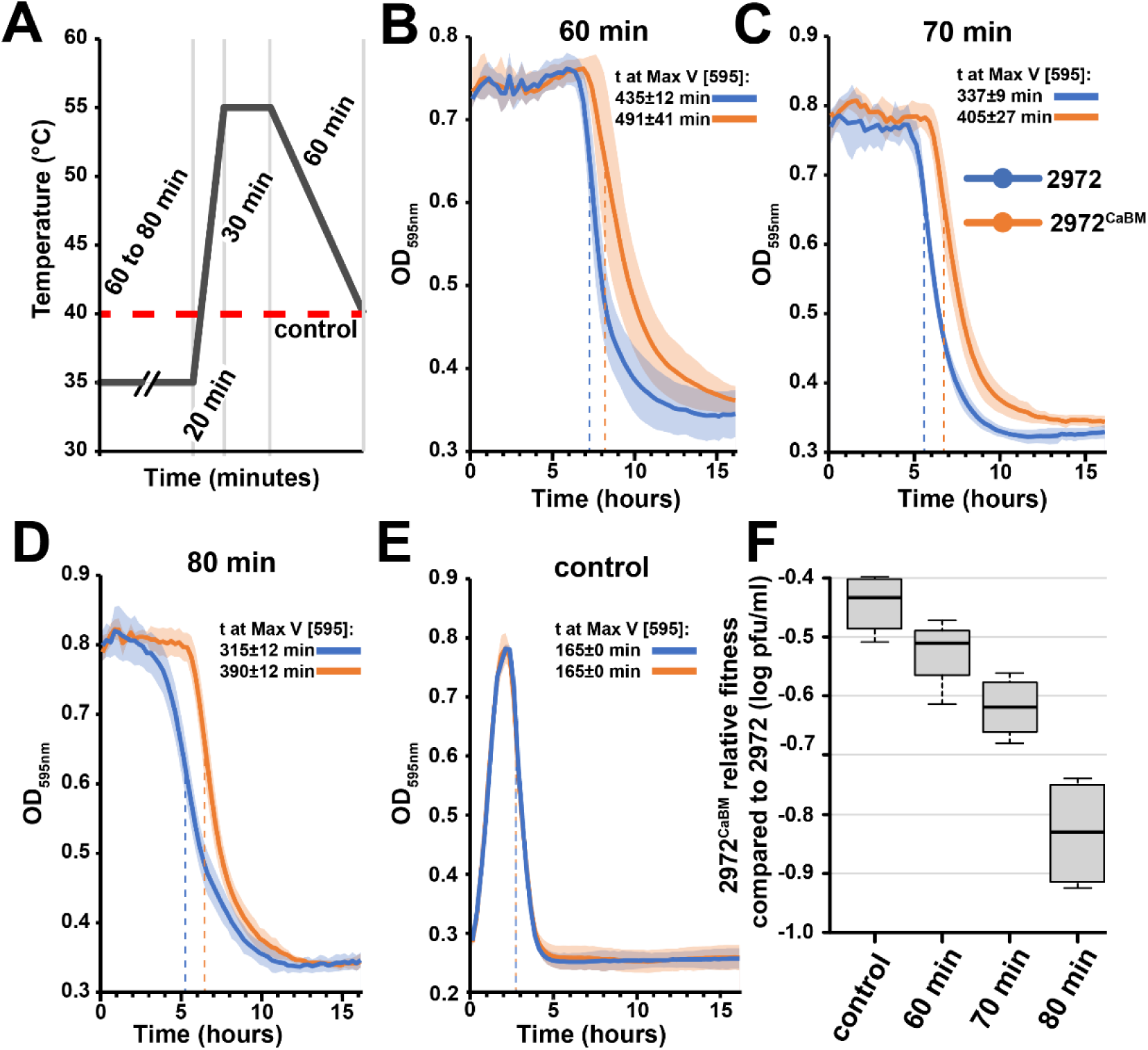
Lytic activity of phages 2972 and 2972^CaBM^ under thermal conditions that are encountered during Emmental cheese production. (A) *S. thermophilus* DGCC7710 cells early in the exponential phase were infected with phage 2972 or 2972^CaBM^ at a MOI of 0.0001. Immediately after infection, cells were incubated in a programmable water bath with a temperature profile used to produce Emmental. The first incubation step at 35 °C was tested for different time periods: **(B)** 60 min early in the exponential phase, **(C)** 70 min, or **(D)** 80 min. After completion of the thermal cycle, the cells were transferred to a microplate reader and turbidity was measured every 15 min over a 16-h period. **(E)** A control was performed by incubating the infected cells at a constant temperature of 40 °C. **(F)** Direct competition assay for relative fitness between phages 2972 and 2972^CaBM^ under the thermal conditions used to produce pressed cooked cheese. Equal amounts of phages 2972 and 2972^CaBM^ were competed against each other at a MOI of 0.0001 and under incubation conditions used in (B) through (E). The titer of phage 2972^CaBM^ after an amplification step was determined on strain *S. thermophilus* DGCC7710 BIM250 (resistant to phage 2972 but not to 2972^CaBM^) and compared with the titer of the total phage population obtained on strain DGCC7710.

The fitness cost on phage 2972^CaBM^ was consistently higher than that observed at a constant temperature of 40 °C, but not for a 60-min incubation step at 30 °C (p-value = 0.1054 for 60 min, p-value = 0.0032 for 70 min, and p-value = 0.0016 for 80 min).

### Phages with a functional CaBM are selected when incubated at high temperature during cheese-making conditions

To further validate the selection of the CaBM under various thermal conditions, we generated a phage mutant with its CaBM inactivated through a single nucleotide substitution (2972^D219A^). Then, we investigated whether the reversion to a functional CaBM could be favored under cheese-making conditions. Specifically, we altered the D219 position, shown to be essential for heat stability, by introducing an alanine through a single nucleotide change in the second codon position (A to C). After subjecting the phage population to experimental evolution (30 transfers) under different thermal conditions (Figure 8A and 8B), we quantified the frequency of the C to A reversion in the CaBM using Illumina MiSeq sequencing. Notably, we did not observe any significant reversion towards a functional CaBM when the phage mutant population was evolved at 37°C. The SNP frequency remained below the detection limit, with a mean frequency of 0.000767±0.000089 at 37°C compared to 0.00125±0.00077 for the control (see Figure 8.C). However, a significant enrichment in SNP frequency leading to a functional CaBM was observed when the phage mutant population was evolved under thermal conditions involving higher temperatures, such as 46°C (0.148±0.252), heat shock (0.0124±0.0184), and cheese-making conditions (0.0142±0.00736).

**Figure 8.**
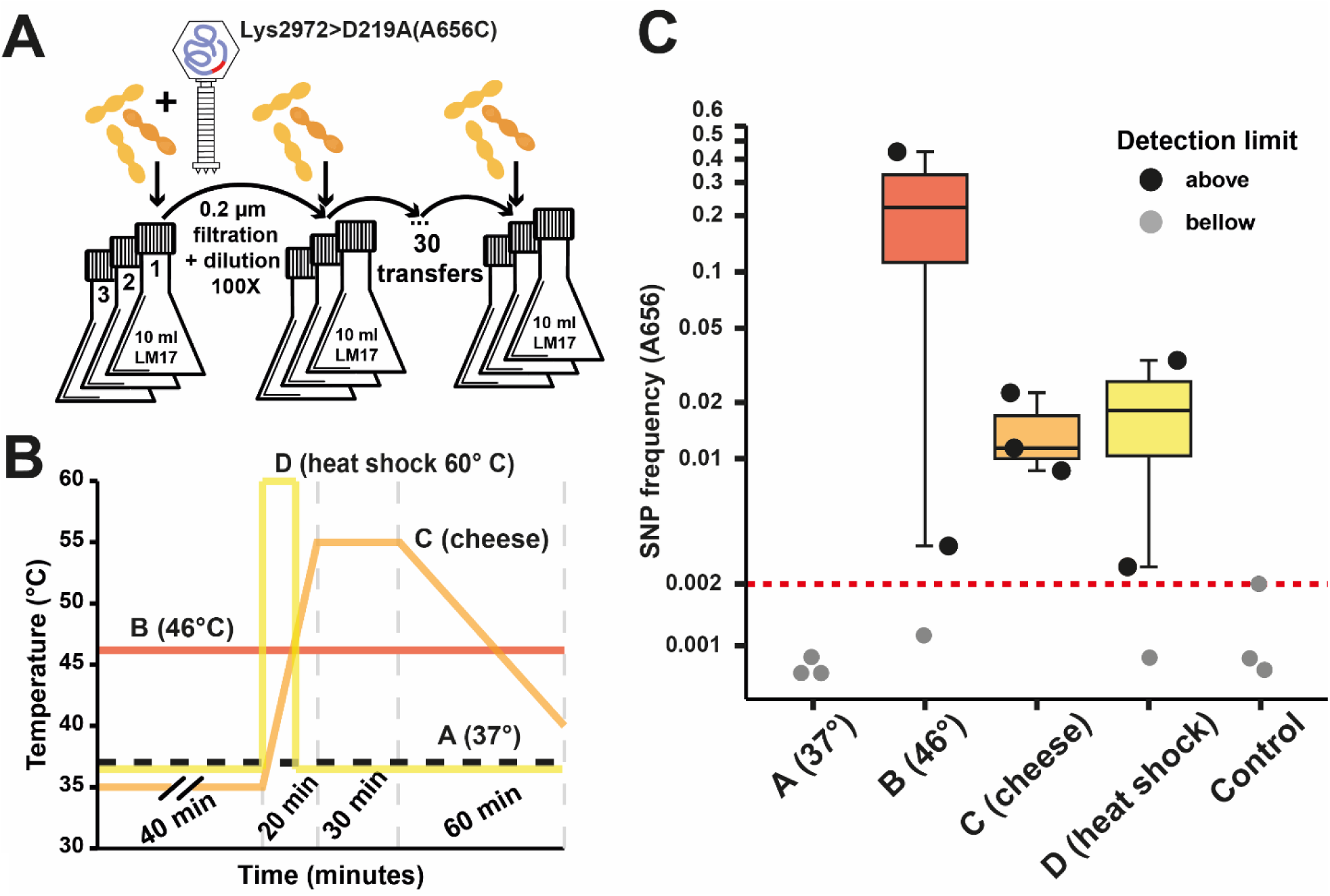
Quantifying the functional reversion of an inactivated CaBM during an experimental evolution assay. A) A phage CaBM mutant with a D219A substitution was created by changing the arginine present in the second position of the domain to a cysteine. The phage mutant was then serially transferred with its host for a total of thirty transfers, each conducted with a final volume of 10 ml and a hundred-fold dilution rate. n=3. **B)** Throughout each of these transfers, phage-infected cells were exposed to a range of thermal conditions, including temperatures of 37°C and 46°C, as well as a 15-minute heat shock at 60°C. Additionally, infected cells were also exposed to thermal conditions that mimic those encountered during the production of pressed cooked cheese. **C)** After the thirty transfers, targeted sequencing of the CaBM at the population level was performed with Illumina MiSeq to measure the SNP frequency (C to A reversion) under the different incubation conditions. The detection limit was set based on the initial SNP frequency found in the phage population before the transfers.

## Discussion

The diversity of endolysins is striking, even among phages that infect the same bacterial species. For example, a total of 26 structural endolysin types have been identified among 200 phages that infect *Mycobacterium smegmatis* ^36^. Similarly, 11 types of endolysins were found in 253 lactococcal phage genomes and each type was associated with a particular phage group ^17^. In this study, we investigated the diversity of endolysins in phages that infect *S. thermophilus*. Surprisingly, we found that endolysin diversity was much lower as only five types were identified in 195 phage genomes.

There are several factors that may have contributed to the low endolysin diversity observed in *S. thermophilus* phages. The first is the limited genetic diversity of strains that were infected with these streptococcal phages. *S. thermophilus* strains are known for their low genetic diversity, as this species is thought to have emerged relatively recently from a commensal ancestor in the Salivarius group ^37,38^. In comparison, the genetic diversity within *L. lactis* is much higher, and up until recently it was divided into four subspecies. Numerous strains of *L. lactis* and the newly created *Lactococcus cremoris* species, are currently used by the dairy industry ^39^. These strains were isolated from various sources such as plants, animals, and raw milk ^40^. Consistent with this lactococcal strain diversity, we recently showed, that the diversity of lactococcal phage endolysins resulted from specific adaptations to the host at the strain level ^17^. Therefore, the significant difference in endolysin diversity between phages that infect these two dairy bacterial species may be due to variations in host diversity, as lower host diversity leads to lower endolysin diversity and vice versa.

Similarly, the limited genetic diversity of *S. thermophilus* strains may affect the diversity of phages that can infect this species ^41^. Indeed, lactococcal phages are currently divided into at least 10 groups ^42^, while *S. thermophilus* phages are classified into 5 groups. Moreover, in our study, it was observed that 62% (120 out of 195) of the analyzed *S. thermophilus*-infecting siphophage genomes belonged to the *Moineauvirus* genus, while 38% (46 out of 195) belonged to the *Brussowvirus* genus. Although these two streptococcal phage groups are by far the most abundant and implicated in milk fermentation failures ^41,43–46^, three other distinct phage groups (*Vansinderenvirus*, *987*, and *P738*) have been recently isolated ^47–49^. Furthermore, *P738*-like phages have distinctive genetic features compared to other dairy streptococcal phage groups and are thought to have evolved separately from them ^49,50^. This unique phage group was the only one that did not possess a type A endolysin, which were otherwise detected in 91% of the genomes examined and in the four other phage groups.

We then hypothesized that type A endolysins had distinctive features to be so prevalent in various streptococcal phages. Structural analysis of the phage 2972 endolysin provided valuable insights into type A endolysins. The analysis not only provided crystallographic data for the amidase_5 and zoocin_A domains, but also revealed the presence of a calcium-binding site (CaBM) in the structure of the endolysin. Calcium is also known to increase the thermal stability of several enzymes including proteases such as trypsin and proteinase K ^30^. Therefore, we investigated the effect of calcium on the thermal stability of Lys2972. Our results showed that the presence of calcium effectively improved the thermal stability of the enzyme and that this effect was not observed in a mutated protein with an inactivated CaBM.

Flexibility in the substrate-binding region is critical for effective substrate identification and binding, as it facilitates mechanisms such as induced fit and conformational selection ^51,52^. The presence of calcium was previously shown to alter the flexibility of the substrate-binding region of proteinase K, thereby affecting the enzyme’s affinity to some extent ^53–55^. Similarly, the presence of calcium may also impact the flexibility of the substrate-binding region of Lys2972, leading to slight variations in enzyme activity. Indeed, our observations confirm that the addition of calcium causes an increase, albeit modest, in enzyme activity, whereas inactivation of CaBM or addition of EDTA resulted in a small decrease.

To gain a deeper understanding of the biological function of CaBM in the lytic cycle of phage 2972, we used the innate CRISPR-Cas system of *S. thermophilus* to directly inactivate the CaBM in the endolysin of phage 2972 by introducing alanine substitutions into the three amino acids known to coordinate calcium. It is noteworthy that despite the mutation of CaBM, replication of the phage was not significantly affected at 37 °C. This is best evidenced by the ability of the mutant phage to produce lytic plaques of comparable size to the wild-type phage, suggesting that CaBM is unlikely to play an important role in either folding or regulation of the enzyme. However, we observed that the fitness of the phage mutants was significantly more impaired when the replication temperature was increased. In addition, the phage mutant was unable to resume cell lysis after a 15-minute heat shock at 60 °C or to induce lysis of a bacterial cell culture at 55 °C, indicating that CaBM plays a critical role in maintaining the thermal stability of the enzyme and making the lytic cycle of phage 2972 more resilient to temperature fluctuations.

The ability of CaBM to thermally stabilize endolysin in the presence of calcium makes it particularly suitable for phages that infect bacteria such as *S. thermophilus*. These bacteria grow in milk, which typically contains about 10 mM of soluble calcium ^56^. In addition, various hard cheeses are made using elevated temperatures. The ability of *S. thermophilus* strains to withstand high processing temperatures has made it an essential ingredient in the production of Swiss cheeses (such as Emmental and Gruyère) and some Italian cheeses (such as Grana Padano and Parmigiano Reggiano), which typically require cooking temperatures in the range of 50 to 58 °C ^33^. The fact that CaBM provides a selective advantage under conditions that simulate a cheese-making process reinforces that thermal stability plays a crucial role in the phage infection of this bacterial species.

In summary, the overrepresentation of type A endolysins in phages that infect *S. thermophilus* is likely due to (1) the low genetic diversity of the host species, which may have favored the use of a single enzyme type; (2) the unique physicochemical properties such as thermostability; and (3) the cheese-making process, which acts as a strong barrier for the selection of adapted phages and their enzymes.

## Methods

### Bacterial strains, phages, and growth conditions

The biological materials and plasmids used in this study are listed in Supplementary Table 1. *S. thermophilus* DGCC7710 and phage 2972 was obtained from the Félix d’Hérelle Reference Center for Bacterial Viruses (www.phage.ulaval.ca). *S. thermophilus* strains were grown without agitation at 42 °C in M17 broth supplemented with 0.5% lactose (LM17) and 1.0% agar (for solid media). When needed, chloramphenicol was added to the media at final concentration of 5 μg/mL (Cm 5). *Escherichia coli* strains were grown at 37 °C with agitation (220 rpm) in LB or plated on LB with agar (LBA). If necessary, LB was supplemented with kanamycin sulfate (30 μg/mL) or chloramphenicol (25 μg/mL for LBA plates and 50 μg/mL for LB). For phage infection, 10 mM CaCl_2_ was added to the M17 media, and the double-layer plaque assays were performed as previously described ^57^.

### Plasmid construction

Plasmids were purified from overnight bacterial cultures using a QIAprep Spin Miniprep Kit (Qiagen). Polymerase chain reactions (PCR) were performed with Taq polymerase (Feldan) for screening purposes or Q5 high-fidelity DNA polymerase (New England Biolabs) for cloning and site-directed mutagenesis. Gibson assembly master mix was prepared as previously described ^58^ and restriction enzymes were purchased from New England Biolabs. Primers used in this study are listed in Supplementary Table 1.

### Phylogenetic analysis and gene conservation of endolysins in virulent phages that infect *S. thermophilus*

A total of 195 phage genomes that were annotated to infect *S. thermophilus* were retrieved from NCBI (02/26/2023). Then, they were analyzed for the presence of endolysin genes and deduced amino acid sequences. Group IA2 introns present in some of endolysin genes were manually removed from group A endolysin sequences. MUSCLE (v3.8.425) was used to perform the multiple sequence alignment with a maximum of 8 iterations ^59^. The phylogenetic tree was generated using Neighbor-Joining and a Jukes-Cantor genetic distance model. The visualization and re-rooting of the tree was done with Interactive Tree Of Life (iTOL) v5 ^60^. The domains were annotated with BLASTP, HHPred, and AlphaFold predictions ^61–63^. The gene conservations were predicted by clustering the genes with cd-hit-est at 80% identity (https://www.ncbi.nlm.nih.gov/pmc/articles/PMC3516142/). The *dn/ds* ratios were calculated by calling single nucleotide polymorphisms on the MUSCLE alignments with freebayes and SNPeffect (https://www.ncbi.nlm.nih.gov/pmc/articles/PMC3245173/).

### Cloning, expression, and purification of Lys2972 and its mutant derivatives

Because of the presence of a group IA2 intron in the gene encoding Lys2972, *lys2972* was amplified from phage 2972 genomic DNA in two fragments with primer pairs Lys2972_F1_Fw / Lys2972_F1_Rv and Lys2972_F2_Fw / Lys2972_F2_Rv. The *lys2972* fragments were then cloned into a pET28a expression vector using Gibson assembly. Primers with complementary ends (35-bp) were designed to bind either to the opposite *lys2972* fragment or to a linearized pET28a vector. The linearized vector was obtained by PCR using the primer pairs pET28a_Fw/pET28a_rv and a BamHI-digested pET28a plasmid as a template. The subsequent plasmid, pLys297228a, was used as a template for site-directed mutagenesis to either introduce alanine substitutions into the Lys2972 CaBM and catalytic site or to generate a CBD-deficient variant. To accomplish this, the Q5® Site-Directed Mutagenesis Kit (NEB) was used with primer pairs designed using the manufacturer’s online software (nebasechanger.neb.com). All constructs were confirmed by sequencing.

For protein purification, the cells were initially cultured overnight in 50 mL of TB media and 5 ml of culture were transferred to a medium volume of 800 mL to achieve an optical density (OD_600nm_) of ∼0.6 before the cells were induced with 1 mM isopropyl-beta-D-1-thiogalactopyranoside (IPTG) and cultured for an additional 16 hours at 18°C. The cell pellets were harvested and suspended in 45 mL of lysis buffer containing 50 mM Tris-HCl pH 7.5, 300 mM NaCl, 5% glycerol, 5 mM imidazole, and 1 mM PMSF. After incubation with DNase (5 ug/ml) and lysosome (2 mg/ml), the cells were lysed using sonication. The resulting lysate was subjected to ultracentrifugation, and the supernatant was then subjected to nickel affinity chromatography. The process involved multiple washes with a gradient imidazole-containing buffer, ranging from 5 mM to 20 mM. Proteins were subsequently eluted using a buffer containing 200 mM imidazole. The eluted protein was applied onto a Superdex 200 Increase (10/300GL) column for buffer exchange. The buffer used for this step consisted of 20mM Tris-HCl pH 7.3, 150 mM NaCl, and 1% glycerol. The pooled fractions were concentrated to 15-20 mg/ml before crystallization setup.

### Crystallization, data collection, and structure determination

The crystallization of Lys2972 was performed at room temperature via the micro-batch-under-oil approach, where the protein and reservoir solution were mixed at a 1:1 ratio (v/v). Lys2972 crystals were obtained using the reservoir solution containing 0.2 M LiCl, 0.1 M NaAc pH 5.0, 20% PEG-6000. The crystals were cryoprotected by the reservoir solution supplemented with 15% ethylene glycol before data collection at the LRL-CAT (31-ID) beamline at Advanced Photon Source, Argonne National Laboratory. Data processing and scaling were performed with XDS ^64^. Lys2972 crystals were in the C2 space group with unit cell parameters of a = 83.2, b = 64.8, c = 55.4 Å, β = 122.7°. The structure was determined by Phaser-MR ^65^ using the templates predicted by Robetta ^66^ followed by multiple cycles of ARP/wARP model building ^67^. To get the final structures, multiple cycles of refinement using Refmac5 ^68^ in CCP4 ^69^ followed by manual model rebuilding with Coot ^70^ were carried out. The stereochemistry of the final model has been analyzed with PROCHECK^71^. Data collection and refinement statistics are shown in Supplementary Table 2.

### Evaluation of the activity of Lys2972 and derivatives on *S. thermophilus* cells

The lytic activity of Lys2972 and its mutant derivatives was evaluated by monitoring the reduction in turbidity of a solution containing *S. thermophilus* DGCC7710 cells at an OD_600nm_ of 0.4. Bacterial cells were resuspended in lysis buffer (20 mM Tris-HCL, 200 mM NaCl, pH 7.0) and mixed in a 96-well microplate with purified endolysin diluted to the desired concentration in endolysin buffer (20 mM Tris pH 7.3, 150 mM NaCl). CaCl_2_ was added to both the endolysin and lysis buffers as needed. The decrease in turbidity was measured at 37 °C using a microplate reader (BioTek Synergy HTX Multimode Reader).

### Evaluation the heat stability and lytic activity of Lys2972 and its CaBM mutants at different temperatures

To conduct thermal shift assays, solutions of Lys2972 and CaBM mutants (each at a concentration of 30 µM) were prepared with or without 10 mM calcium. Next, 20 μl of the prepared protein were mixed with 5 μl of 500-fold diluted JBS Thermofluor Dye (Jena Bioscience), prepared in the same buffer. After an initial incubation at 4 °C for 2 minutes, samples were gradually heated at a rate of 0.5 °C/min, ending at 95 °C, using a CFX96 Touch RealTime PCR Detection System (Bio-Rad). The FRET channel was used to collect data during the heating process, and the results were analyzed using the Bio-Rad CFX Maestro software.

The lytic activity of Lys2972 and Lys2972^CaBM^ was also tested in the presence or absence of calcium and after an incubation of 15 minutes at different temperatures. Solutions of the two endolysins were prepared at a final concentration of 30 µM with endolysin buffer with or without 10 mM calcium. Each solution was then distributed into separate 500 µL PCR tubes in 100 µL aliquots and exposed to different temperatures using a thermal cycler. After thermal exposure, the tubes were centrifuged at 13000 rpm for 2.5 minutes and the resulting supernatant was diluted tenfold before being tested for lytic activity using the same methodology as previously described.

To evaluate the lytic activity of Lys2972 and Lys2972^CaBM^ at different temperatures, 100 ul of *S. thermophilus* DGCC7710 cells in the exponential phase (resuspended in lysis buffer) and 200 ul of endolysin (4 µM) were pre-incubated in 500 ul PCR tubes. The pre-incubation was done in a water bath at temperatures of 37 °C, 45 °C, 55 °C, or 60 °C, with 10 mM CaCl_2_ as required. After 15 minutes of pre-incubation, 100 ul of endolysin was added to the tubes containing the bacterial cells, and the decrease in turbidity was measured 15 minutes later using a DeNovix DS-11 Spectrophotometer.

### Genome editing of phage 2972

CRISPR-Cas-based genome editing was used to inactivate Lys2972 CaBM in the phage 2972. To achieve this, a natural CRISPR BIM (Bacteriophage Insensitive Mutant) targeting the C-terminal part of Lys2972 was isolated as previously described ^72^ and designated as *S. thermophilus* DGCC 7710 BIM-Lys2972 (Supplementary Table 1). Subsequently, repair templates were created by introducing mutations into the Lys2972 CaBM motif. Specifically, the codons encoding residues N208, D216, and D219 were replaced with codons encoding alanine (phage mutant 2972^CaBM^, or alternatively, only D219 was replaced with alanine (phage mutant 2972^D219A^). The mutated CaBM was flanked on each side by approximately 250 bp of phage genomic homologous regions and an additional synonymous mutation in the adjacent protospacer motif (PAM) was inserted to prevent CRISPR-Cas interference with the repair template without changing. The gene fragment (gBlock) was synthesized with complementary ends (Integrated DNA Technologies, Inc.) for Gibson assembly at the XbaI restriction site of plasmid pNZ123 ^73^. Recombinant phages were isolated from plaques after the infection of *S. thermophilus* DGCC 7710 BIM-Lys2972 strain transformed with the repair template plasmid. Modified phages with the correct mutations were screened with primer pairs 2972_CaCl2_Fw and 2972_CaCl2_Rv. Subsequently, the genome of one of the phage mutants, designated as 2972^CaBM^, was fully sequenced.

### Phage DNA sequencing and analysis

Phage genomic DNA was extracted following the procedures outlined elsewhere ^74^. Libraries were prepared using the Nextera XT DNA library preparation kit (Illumina), adhering to the manufacturer’s instructions. Sequencing was carried out on a MiSeq system, utilizing a MiSeq reagent kit v2 (Illumina). Subsequently, the reads underwent cleaning through Trimmomatic v0.36 ^75^ and were assembled to obtain circular complete sequences, utilizing Ray v3.0.1 ^76^ and SPAdes v3.13 ^77^.

### Quantification of phage Lys2972 and CaBM mutant lytic activity, plaque size, and fitness at different temperatures

To investigate the effects of CaBM inactivation on phage 2972 lytic activity, *S. thermophilus* DGCC7710 cells were infected with either phage 2972 or 2972^CaBM^ at a MOI of 0.1 at an OD_600nm_ of 0.2. After infection, 50-µl aliquots were transferred into 200-µl PCR tubes and exposed to different temperatures in a thermal cycler. After 16 hours, the absorbance of the cell cultures was measured using a DeNovix DS-11 spectrophotometer.

Lysis of *S. thermophilus* DGCC7710 cells during infection with phage 2972 or 2972^CaBM^ was also monitored at different temperatures. To achieve this, 10-ml cultures were grown in glass tubes until they reached an OD_600nm_ of 0.). Cells were then infected with phage 2972 or 2972^CaBM^ at a MOI of 0.01 and incubated in water baths set at 37 °C, 42 °C, or 46 °C. Absorbance was monitored every 30 minutes for 3 hours using a SPECTRONIC 20^+^ spectrophotometer (Spectronic Instruments).

To analyze the size of lytic plaques generated by phage 2972 and 2972^CaBM^, phage lysates were diluted to produce 50 to 100 lytic plaques per plate. The plates were then incubated at 37 °C, 42 °C, or 46 °C. The surface area of the lytic plaques was determined by image analysis as previously described ^17^.

To evaluate the fitness costs associated with CaBM inactivation, phages 2972 and 2972^CaBM^ were first co-amplified at different temperatures. *S. thermophilus* DGCC7710 cells at an OD_600nm_ of 0.2 were infected with both phage 2972 and 2972^CaBM^ at an initial MOI of 0.0001 and incubated overnight at 37 °C, 42 °C, or 46 °C in different water baths. The fitness cost associated with CaBM inactivation was determined by calculating the difference between the titer of phage 2972^CaBM^ and the titer of the total phage population. To determine the number of 2972^CaBM^ particles in the mixed population, a plaque assay was performed using the strain *S. thermophilus* DGCC7710 BIM-Lys2972. This strain contains a spacer that specifically targets *lys2972*, resulting in a significant 5.0 ± 0.1 log pfu/ml reduction of the phage 2972 population. However, this reduction does not apply to 2972^CaBM^, as its PAM was modified to prevent any interference with the CRISPR-Cas system during genome editing. A parallel titration was performed with strain *S. thermophilus* DGCC7710 to determine the titer of the entire phage population.

### Quantification of phage Lys2972 and CaBM mutant lytic activity after heat shock or under conditions that mimic cheese production

To investigate the effects of CaBM inactivation on heat shock resistance, *S. thermophilus* DGCC7710 cells were cultured in glass tubes until they reached the early exponential phase. The cells were then infected with either phage 2972 or 2972^CaBM^ at a MOI of 0.1. Following infection, the cultures were subjected to heat treatment for 15 minutes by immersing the tubes in a 60 °C water bath, either immediately or at 10-minute intervals. After the heat shock treatment, the cultures were maintained in a water bath at 42 °C, and the cell turbidity was monitored every 30 minutes for 5.5 hours using a SPECTRONIC 20^+^ spectrophotometer.

To investigate the impact of CaBM inactivation in thermal conditions that replicate the cheese-making process, *S. thermophilus* DGCC7710 cells were cultured in glass tubes until they reached the early exponential phase. The cells were then infected with either phage 2972 or 2972^CaBM^ at a MOI of 0.0001. Subsequently, the tubes were transferred to a programmable water bath that replicated the temperature gradients during cheese production. This involved an initial incubation period at 35 °C for 60 to 80 minutes, followed by a temperature shift to 55 °C over 20 minutes, which was then maintained for an additional 30 minutes. Finally, the temperature was decreased to 40 °C over one hour. After these incubations, the infected cells were transferred to a 96-well plate and incubated at 40 °C in a microplate reader and absorbance was measured every 15 minutes for 16 hours (BioTek Synergy HTX Multimode Reader). Time at maximum rate (t at max V) of cell lysis kinetics was determined using BioTek Gen5 software (v3.12).

The fitness costs associated with CaBM were assessed by co-amplifying phages 2972 and 2972^CaBM^ at a MOI of 0.0001, under the same thermal conditions that were used to replicate the cheese-making process. The fitness cost was determined by calculating the difference between the titer of phage 2972^CaBM^ and the titer of the total phage population using strain *S. thermophilus* DGCC7710 BIM-Lys2972, as described previously.

### Experimental evolution of Lys2972 CaBM

The phage CaBM mutant 2972^D219A^ was successfully amplified in triplicates for a total of 30 transfers. *S. thermophilus* DGCC7710 cells in the exponential growth phase (OD600nm 0.2) were infected with an initial MOI of 0.01. Infected cells were then incubated at different temperatures: 37°C, 46°C, or for an additional period of 40 minutes at 42°C before being subjected to a 15-minute heat shock at 60°C. Additionally, infected cells were also exposed to thermal conditions mimicking those encountered during the production of pressed cooked cheese, 40 minutes post-infection. After an incubation period of 12 hours at 30°C, phage lysates were filtered and diluted a hundred times before starting another round of amplification.

After 30 transfers, phage DNA was extracted, and the CaBM motif was sequenced using Illumina MiSeq V2. Libraries for high-throughput sequencing of PCR products were prepared following the “16S Metagenomic Sequencing Library Preparation” protocol by Illumina, using primers with extensions containing the adapters and the MiSeq Reagent Kit v2 (CaBM_D219A_Fw and CaBM_D219A_Rv, Supplementary Table 1). Adaptors and low-quality bases were trimmed from the raw reads using trimGalore with the default parameters. Subsequently, the filtered reads were mapped to the 200 bp reference using bwa mem ^78^. Over 99.2% of reads mapped in all samples, and the reference was sequenced with an average depth of 10⁶ coverage (Supplementary Figure 10). Single nucleotide polymorphisms (SNPs) were called using freebayes (-F 0.01 -p 12 –pooled- discrete) ^79^. To establish the detection limit, we considered 0.002, which represented the highest SNP frequency observed in the control. Coincidentally, the default Phred quality score of 40 corresponds to the 0.002 technical error rate, serving as the technical error threshold.

## Acknowledgements

The authors would also like to thank Denise Tremblay, Genevieve Rousseau, Alessandra Gonçalves de Melo and Rachel Morin-Pelchat for technical help and suggestions. We thank Amanda Toperoff and Michi Waygood for editorial assistance. F.O. was supported by a post-doctoral fellowship from the Swiss National Science Foundation under grant P400PB_191059. C.M. and X.Z. were supported by graduate scholarships from the Fonds de Recherche du Québec - Nature et Technologies. R.S. and S.M. are thankful to PROTEO for funding through their new initiative program. S.M. also acknowledges funding by the Natural Sciences and Engineering Research Council of Canada (Discovery and Alliance programs). S.M holds a T1 Canada Research Chair in Bacteriophages. V.S. was supported by a post-doctoral fellowship from the Swiss National Science Foundation under grant P500PB_214419. This research used resources of the Advanced Photon Source, a U.S. Department of Energy (DOE) Office of Science User Facility operated for the DOE Office of Science by Argonne National Laboratory under Contract No. DE-AC02-06CH11357. Use of the Lilly Research Laboratories Collaborative Access Team (LRL-CAT) beamline at Sector 31 of the Advanced Photon Source was provided by Eli Lilly Company, which operates the facility.

