## Supplementary Figures for "Dairy practices select for thermostable endolysins in phages"

**Supporting information**

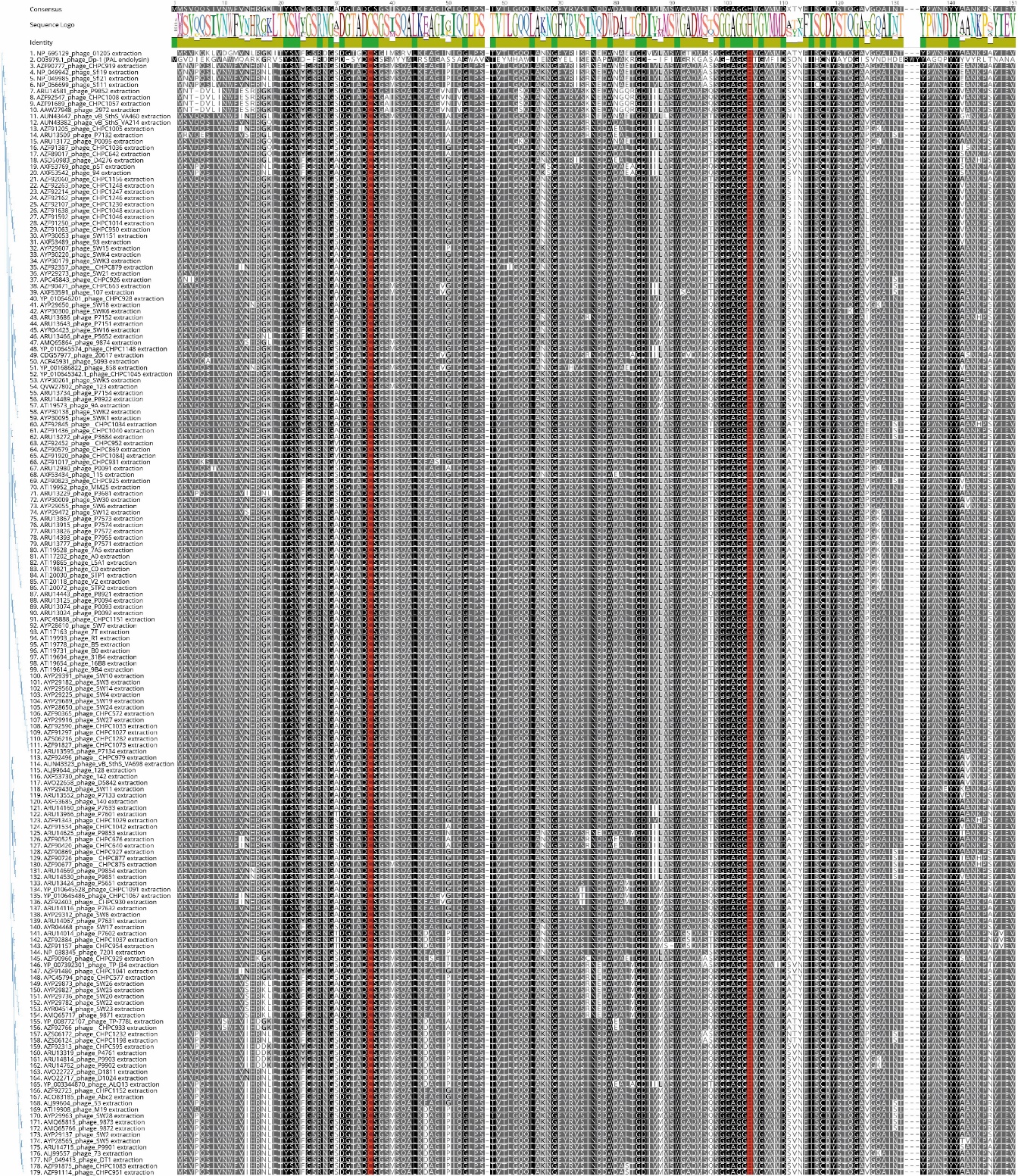

**Supplementary Figure 1. Multiple sequence alignment of the amidase_5 EAD domains of endolysins belonging to cluster A of phages that infect *S. thermophilus* and the PAL endolysin from phage Dp-1 that infects *S. pneumonia*.** Similar residues are colored according to their level of conservation based on BLOSUM62 scores (black: 100% similar, grey: 80 to 99%, light grey: 60 to 79%, white: less than 60%). The identity of all the pairs in the column is indicated on the top (green: 100% identity, green-brown: 30% to 99% identity, red: less than 30% identity). The conserved cysteine and histidine residues that are part of the CHAP active site are highlighted in red. The alignment and figure were generated using the Geneious v11.1.5 software package ^1^.

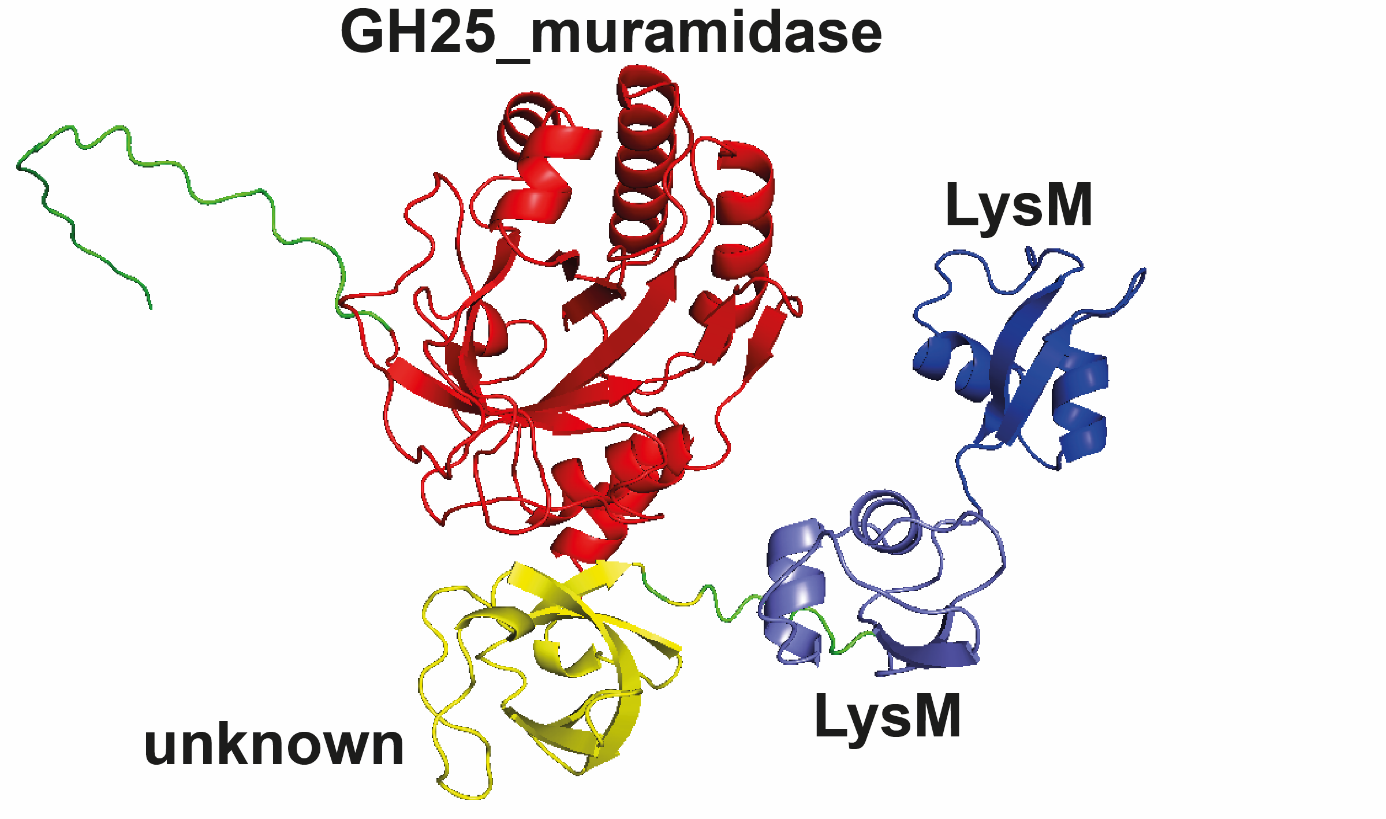

**Supplementary Figure 2. AlphaFold structural prediction of the *S. thermophilus* phage CHPC1109 endolysin (group E).** The tertiary structure of LysCHPC1109 was predicted using ColabFold ^2^. The domains of the endolysin are shown in different colors and named according to HHPred and BLASTP predictions.

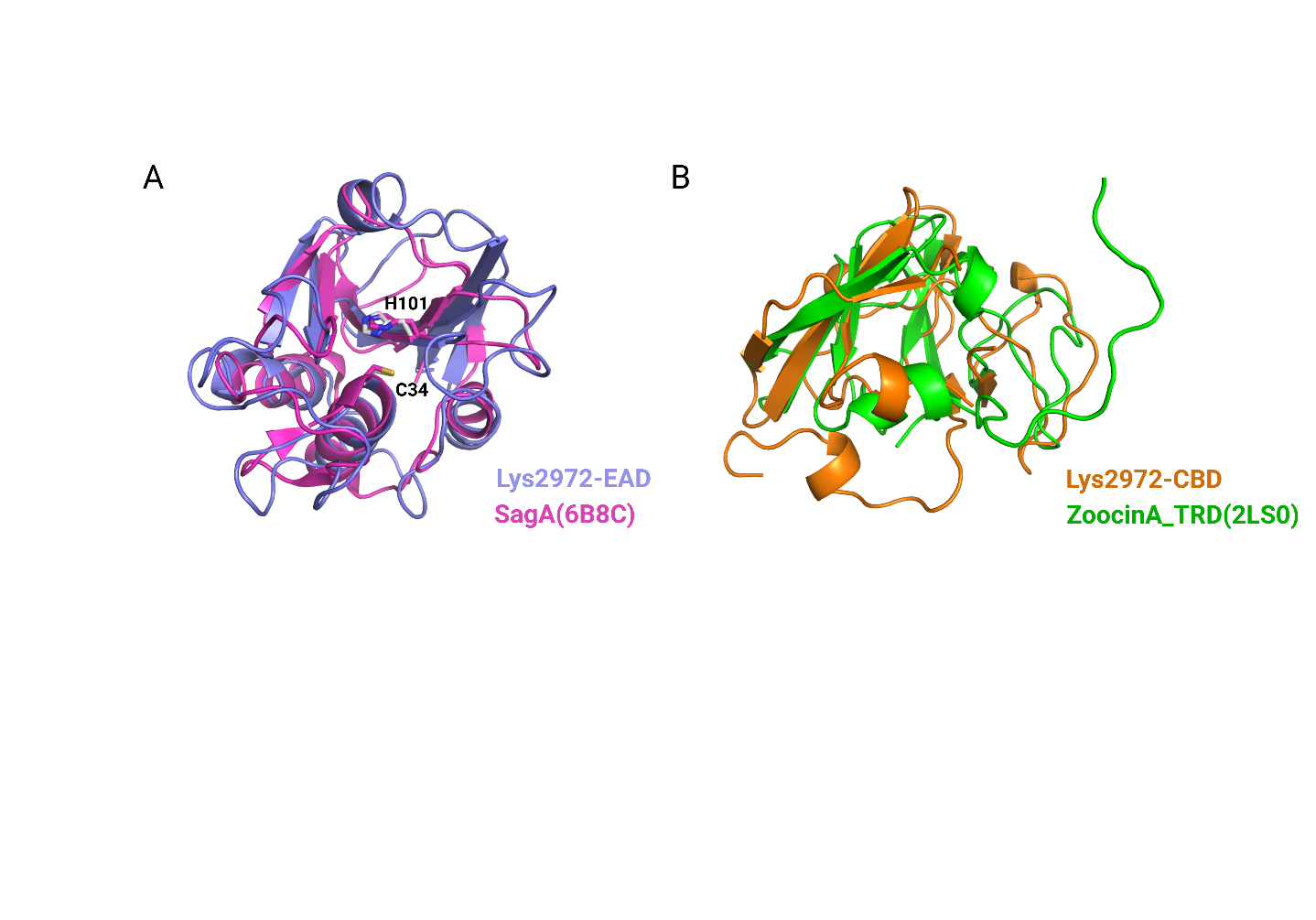

**Supplementary Figure 3. Superposition of Lys2972 domain with its structural homologue.**  A) Superposition of Lys2972 EAD domain (colored in light blue) and SagA (PDB 6B8C, colored in light magenta). The catalytic residues C34 and H101 of Lys2972 are shown in stick models. B) Superposition of Lys2972 CBD domain (colored in orange) and the target recognition domain (TRD) of Zoocin_A (PDB 2LS0, colored in green). The r.m.s.d. values for the aligned Ca atoms are both 2.1 Å for SagA (106 Ca atoms) and Zoocin_A TRD (94 Ca atoms), respectively.

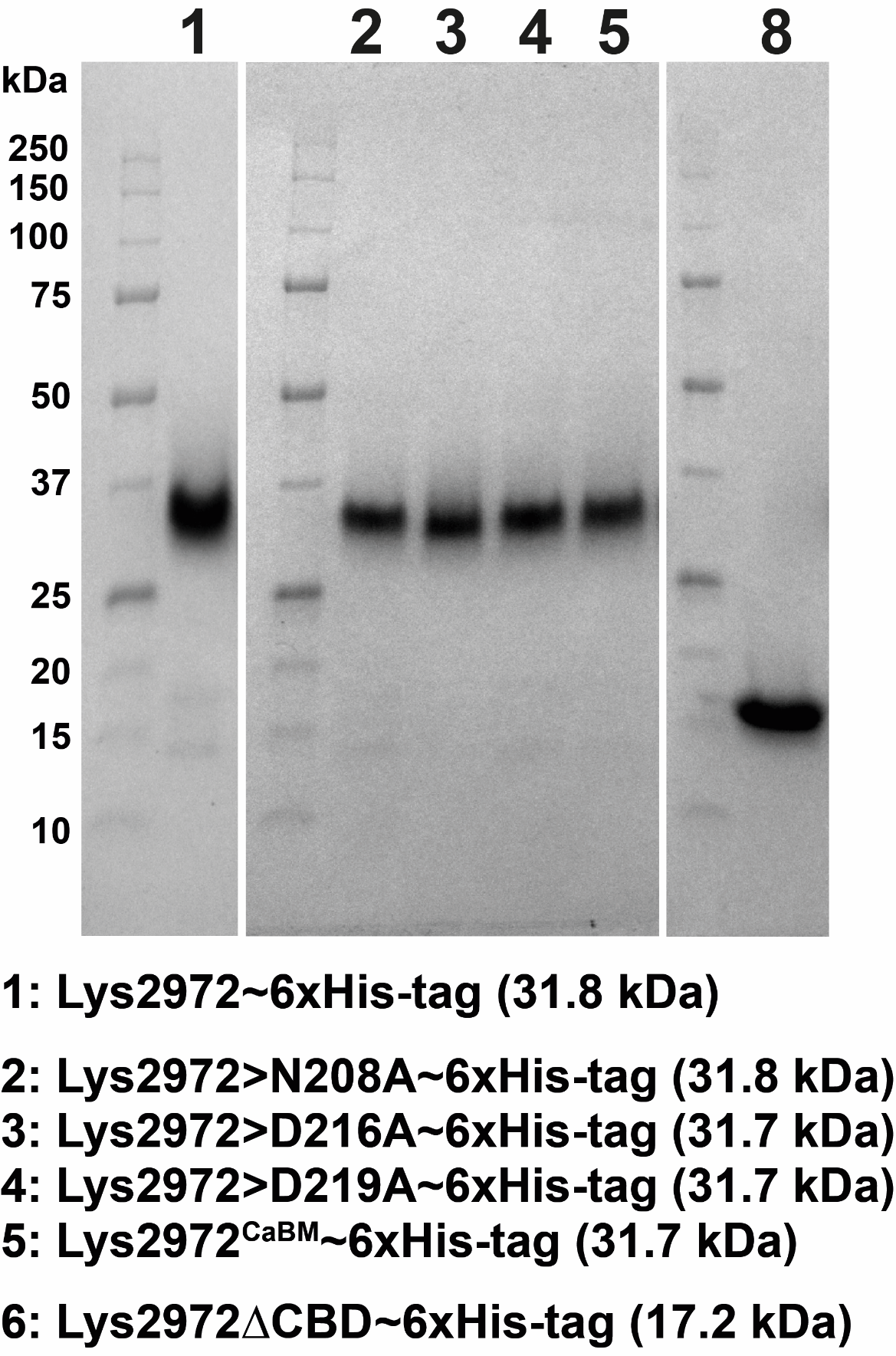

**Supplementary Figure 4. SDS-PAGE gel showing the purified endolysins used in the study**. Proteins were loaded on NuPAGE 4–12% BisTris gels and stained with Coomassie Blue. The expected molecular mass of each purified protein is indicated, and molecular weight markers are shown on the left of each gel.

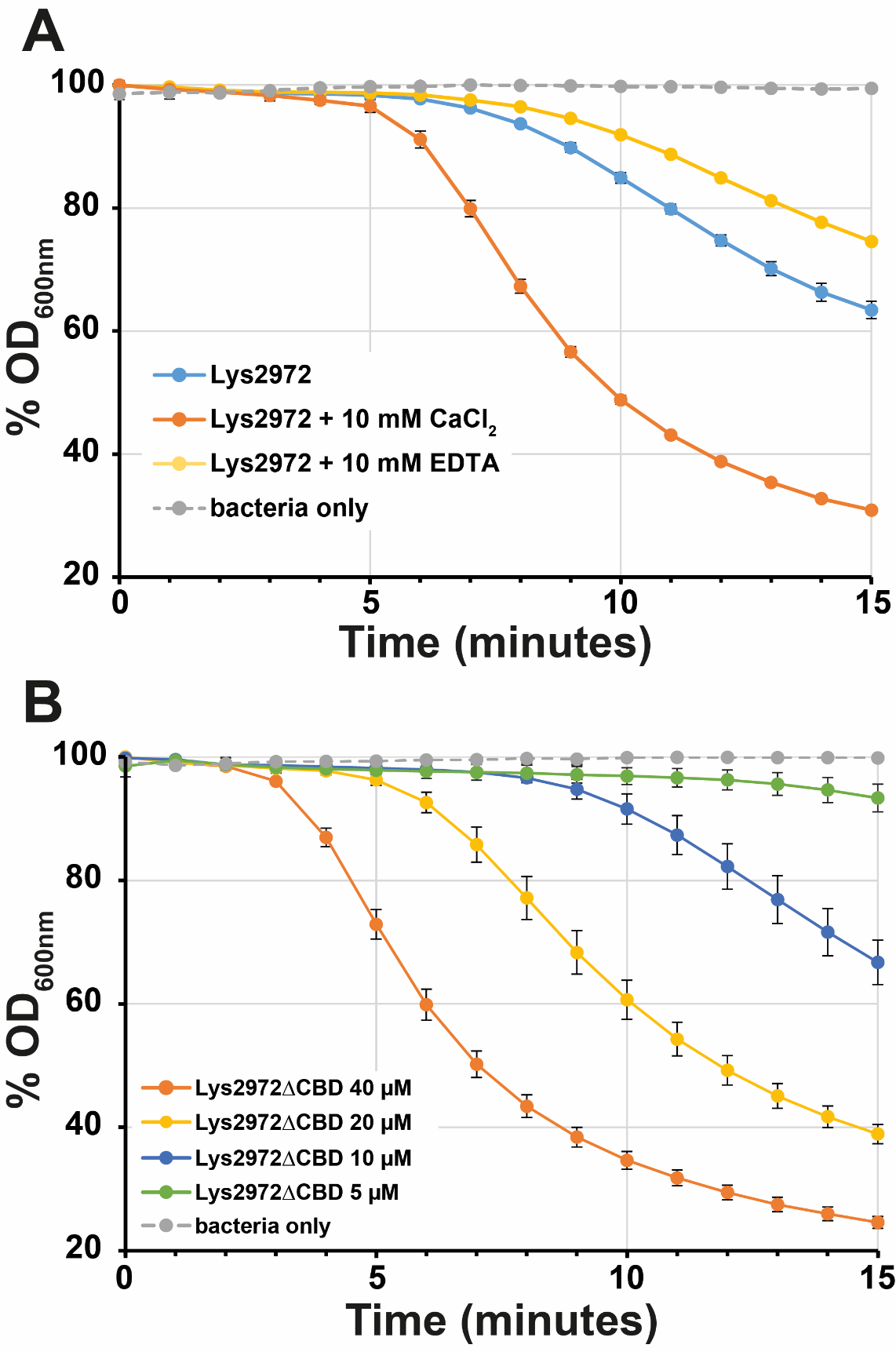

**Supplementary Figure 5. Analysis of Lys2972 enzymatic activity following EDTA addition, CBD deletion, and catalytic site inactivation. A)** The activity of Lys2972 (0.1 µM) was assessed in the presence of either 10 mM CaCl2 or 10 mM EDTA. **B)** Enzymatic activity of Lys2972ΔCBD evaluated at various concentrations. The activity was measured by monitoring the decrease in turbidity of *S. thermophilus* DGCC7710 cells in the exponential growth phase every minute for 15 minutes. Experiments were performed in triplicate.

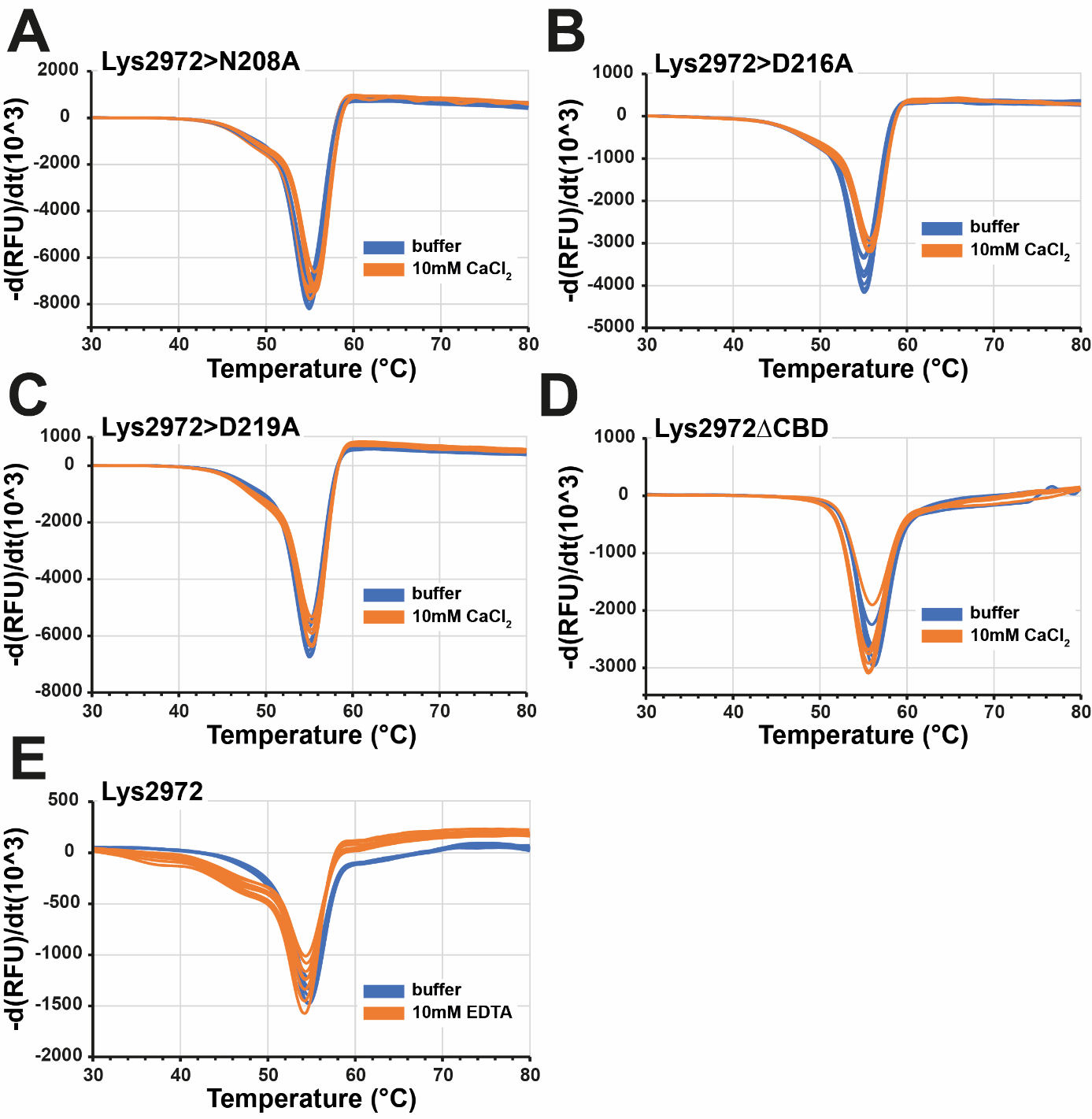

**Supplementary Figure 6. Thermal shift assay of wild-type and mutated Lys2972 containing an inactivated CaBM through single amino acid substitution or deletion of the CBD, or Lys2972 in the presence of EDTA.** The experiments were performed using SYPRO® Orange, and the Tm (lowest point of the curve) was determined by plotting the first derivative of the fluorescence emission against temperature steps of 0.5 °C.

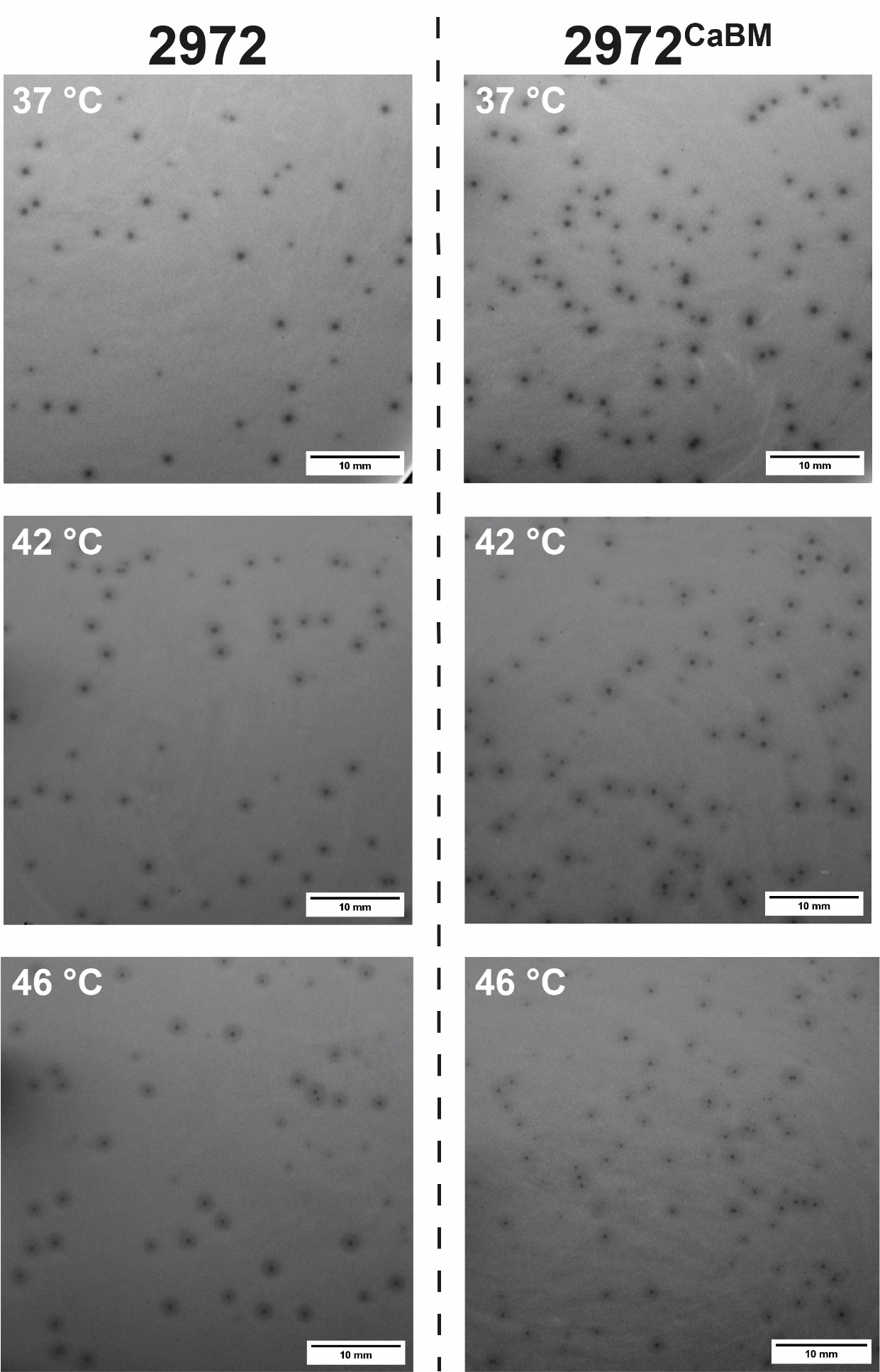

**Supplementary Figure 7. Examples of plaques obtained with lytic phages 2972 and 2972^CaBM^ at different temperatures using *S. thermophilus* DGCC7710 as host strain.** Images of the lytic plaques were obtained at a resolution of 35 pixels per mm.

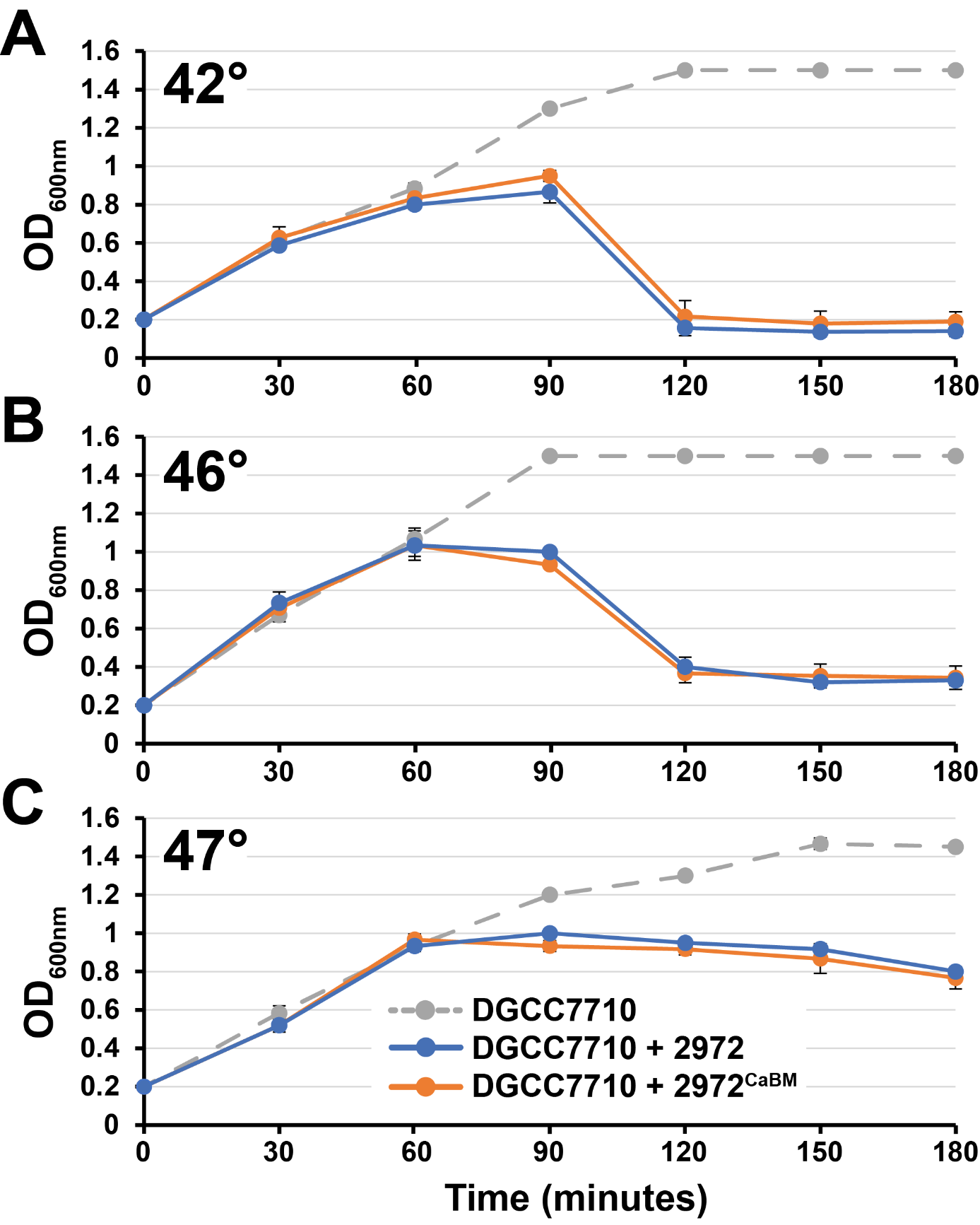

**Supplementary Figure 8. Lytic activity of phage 2972 and 2972^CaBM^ at different incubation temperatures.** *S. thermophilus* DGCC7710 cells were infected in the early exponential phase (OD_600nm_ 0.2) at a MOI of 0.01. After infection, cells were incubated at different temperatures and optical density was measured every 30 minutes. Experiments were performed in triplicate.

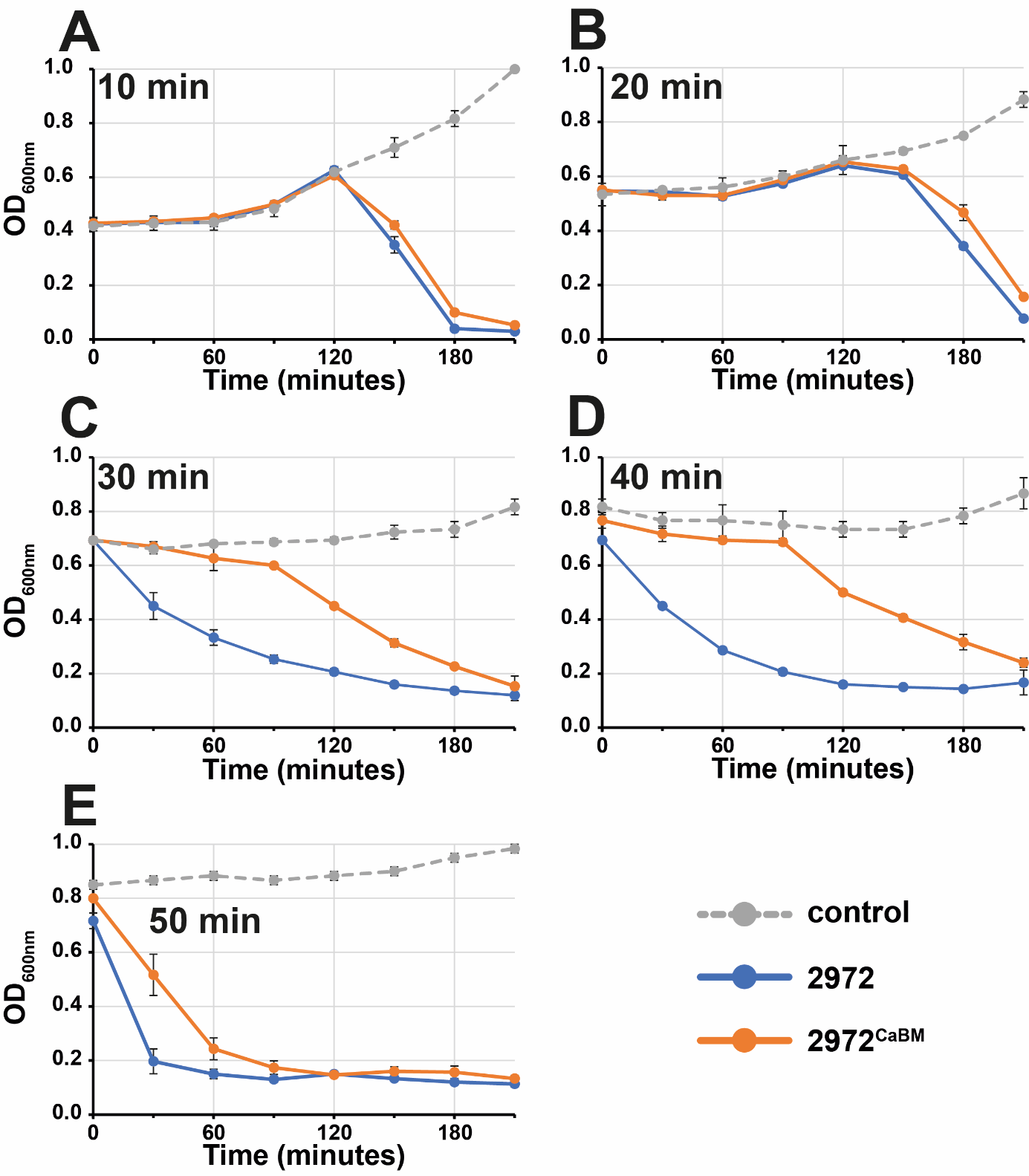

**Supplementary Figure 9. Effects of a heat shock at 55 °C on the lytic activity of phages 2972 and 2972^CaBM^.** *S. thermophilus* DGCC7710 cells in the early exponential phase were infected with phage 2972 or 2972^CaBM^ at a MOI of 0.1. Infected cells were then subjected to heat shock at 55 °C for 15 minutes either (A) immediately or (B-F) at various time points after infection. Optical density was measured every 30 minutes after heat shock. Experiments were performed in triplicate.

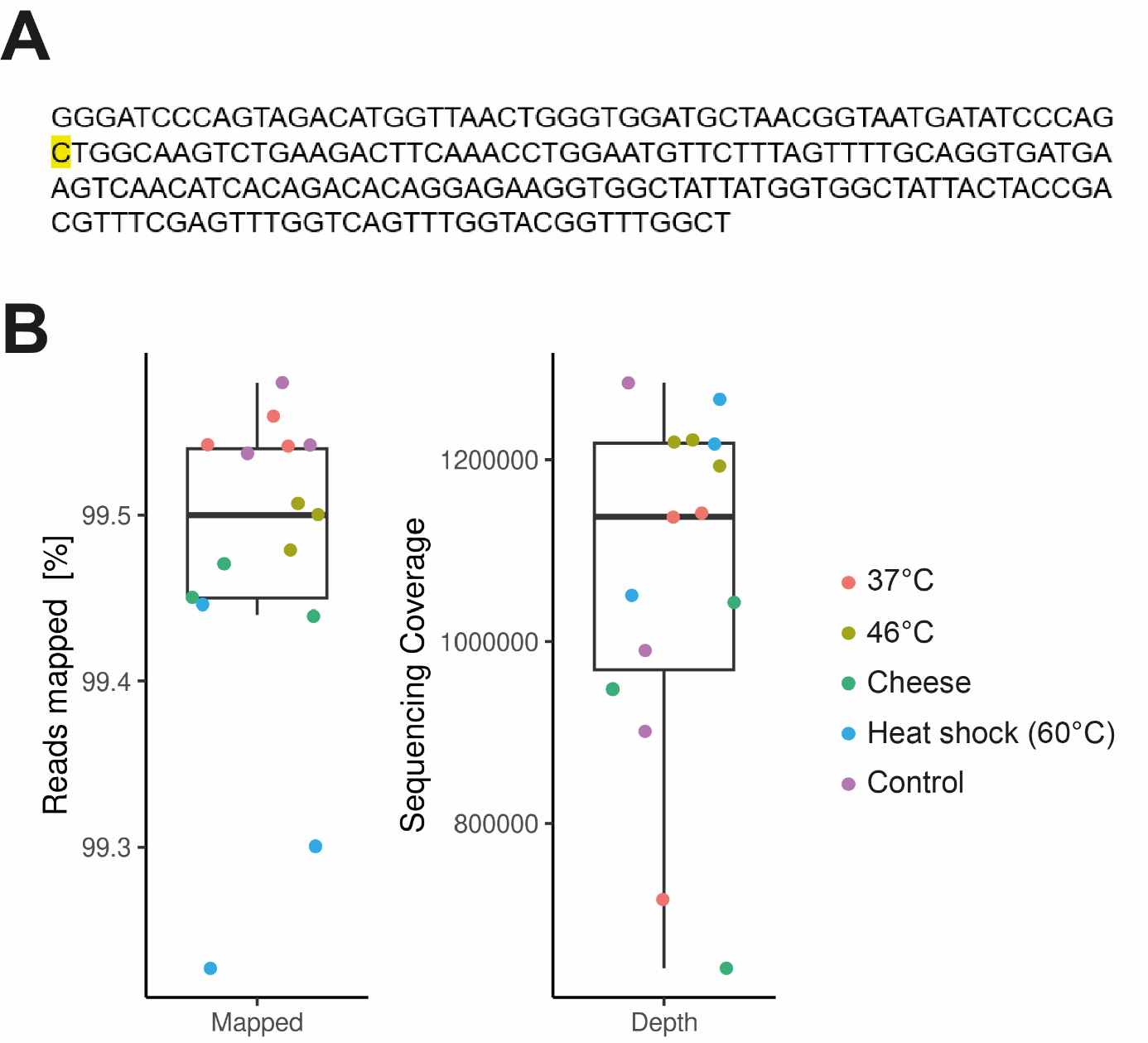

**Supplementary Figure 10. A)** Reference sequence containing the inactivated CaBM of phage 2972D219A in which the D219 position was altered by the introduction of alanine through a single nucleotide substitution (A - > C, highlighted in yellow). **B)** Percentage of reads mapped against the 200 bp CaBM reference sequence of phage 2972D219A and sequencing depth for each experimental condition shown in Figure 8.
