## Supplementary Table 2 for "Dairy practices select for thermostable endolysins in phages"

**Supplementary Table 2:** X-ray data collection and refinement statistics (Values in parentheses are for the highest-resolution shell).

| **Structure** | **Lys2972** |
| --- | --- |
| **Space group** | **C2** |
| *a, b, c* (Å) | 83.2, 64.8, 55.4 |
| α, β, γ (°) | 90, 122.7, 90 |
| wavelength(Å) | 0.9793 |
| resolution (Å) | 47.6-1.27 (1.34-1.27) |
| observed *hkl* | 210090 (31264) |
| unique *hkl* | 63597 (9345) |
| redundancy | 3.3 (3.3) |
| completeness (%) | 97.6 (98.4) |
| R*_meas_* | 0.043 (0.671) |
| CC_1/2_ | 0.999 (0.684) |
| I/(σI) | 13.6 (2.3) |
| Wilson B (Å^2^) | 16.1 |
| R*_work_* (# *hkl*) | 0.169 (60507) |
| R*_free_* (# *hkl*) | 0.191 (3086) |
| B-factors (Å^2^) |  |
| (# atoms) |  |
| Protein | 20.3 (2163) |
| Ligand | 20.7 (7) |
| Water | 31.9 (249) |
| Ramachandran |  |
| favored (%) | 99.6 |
| Generally allowed (%) | 0.4 |
| outlier(%) | 0 |
| rmsd |  |
| bonds (Å) | 0.0153 |
| angles (°) | 1.94 |
| PDB code | 8TW1 |
