## Supplementary Table 1 for "Dairy practices select for thermostable endolysins in phages"

**Supplementary Table 1:** Bacterial strains, bacteriophages, plasmids, and nucleotides.

| **Bacterial strains** | **Characteristics** | | **Origin** |
| --- | --- | --- | --- |
| ***E. coli*** |  | |  |
| BL21(DE3)pLysS | F^-^ *ompT hsdSB* (r_B_ ^-^ m_B_^-^) *dcm*^+^*gal* (DE3) pLysS (Cam^r^ ) | | Stratagene |
| BL21/pLys2972^28a^ | *E. coli* BL21 (DE3) transformed with pLys2972^28a^ | | This study |
| BL21/pLys2972>N208A^28a^ | *E. coli* BL21 (DE3) transformed with pLys2972>N208A^28a^ | | This study |
| BL21/pLys2972>D216A^28a^ | *E. coli* BL21 (DE3) transformed with pLys2972>D216A^28a^ | | This study |
| BL21/pLys2972>D219A^28a^ | *E. coli* BL21 (DE3) transformed with pLys2972>D219A^28a^ | | This study |
| BL21/pLys2972^CaBM 28a^ | *E. coli* BL21 (DE3) transformed with pLys2972^CaBM 28a^ | | This study |
| BL21/pLys2972ΔCBD^28a^ | *E. coli* BL21 (DE3) transformed with pLys2972ΔCBD^28a^ | | This study |
| ***S. thermophilus*** |  | |  |
| DGCC7710 | *S. thermophilus* laboratory strain | | ^1^ |
| DGCC7710 BIM-Lys2972  (SMQ-1490) | *S. thermophilus* DGCC7710 CRISPR BIM (Bacteriophage Insensitive Mutant) targeting the C-terminal part of Lys2972 with spacer sequence TTCCAGGTTTGAAGTCTTCAGACTTGCCAT | | This study |
| DGCC7710 BIM-Lys2972 / pLys2972^CaBM 123^ | DGCC7710 BIM-Lys2972 transformed with pLys2972^CaBM 123^ | | This study |
| DGCC7710 BIM-Lys2972 / pLys2972^D219A 123^ | DGCC7710 BIM-Lys2972 transformed with pLys2972^D219A 123^ | | This study |
| **Phages** | **Characteristics** | | **Origin** |
| 2972 | Phage infecting *S. thermophilus* DGCC7710 | | ^2^ |
| 2972^CaBM^ | Phage 2972 carrying the *Lys2972*>*N208A+D216A+D219A* mutation | | This study |
| 2972^D219A^ | Phage 2972 carrying the *Lys2972*>*D219A* mutation | | This study |
| **Plasmids** | **Characteristics^a^** | | **Origin** |
| pET-28a | Expression vector; Kan^r^ | | Novagen |
| pLys2972^28a^ | pET-28a carrying *Lys2972* | | This study |
| pLys2972>N208A^28a^ | pET-28a carrying *Lys2972>N208A* | | This study |
| pLys2972>D216A^28a^ | pET-28a carrying *Lys2972>D216A* | | This study |
| pLys2972>D219A^28a^ | pET-28a carrying *Lys2972>D219A* | | This study |
| pLys2972^CaBM 28a^ | pET-28a carrying *Lys2972>N208A+D216A+D219A* | | This study |
| pLys2972ΔCBD^28a^ | pET-28a carrying *pLys2972ΔCBD* | | This study |
| pLys2972^CaBM 123^ | pNZ123 having the repair template for the *Lys2972*>*N208A+D216A+D219A* substitution in phage 2972 | | This study |
| pLys2972^D219A 123^ | pNZ123 having the repair template for the *Lys2972*>*D219A* substitution in phage 2972 | | This study |
| pNZ123 | High copy number vector, Cmr, 2.5 kb | | ^3^ |
| **Oligonucleotides** | **Sequence 5’ → 3’** | | **Origin** |
| pET28a_Fw | CACCACCACCACCACCACTGA | | This study |
| pET28a_Rv | CATGGTATATCTCCTTCTTA | | This study |
| Lys2972_F1_Fw | AATTTTGTTTAACTTTAAGAAGGAGATATACCATGAATACAGATGTTTTAATCAATTGG | | This study |
| Lys2972_F1_Rv | TAACCATGTCTACTGGGATCCCATTTTCTCCCCAGTCGAATCCAACGG | | This study |
| Lys2972_F2_Fw | ACTATTTGGCACCCGTTGGATTCGACTGGGGAGAAAATGGGATCCC | | This study |
| Lys2972_F2_Rv | TTAGCAGCCGGATCTCAGTGGTGGTGGTGGTGGTGTTGGTAATAGTTTACCAAATC | | This study |
| Lys2972_N208A_Fw | AGACATGGTTGCATGGGTGGATGC | | This study |
| Lys2972_N208A_Rv | ACTGGGATCCCATTTTCTC | | This study |
| Lys2972_D216A_Fw | TAACGGTAATGCAATTCCAGATGGC | | This study |
| Lys2972_D216A_Rv | GCATCCACCCAGTTAACC | | This study |
| Lys2972_ D219A_Fw | TGATATTCCAGCAGGCAAGTCTGAAG | | This study |
| Lys2972_ D219A_Rv | TTACCGTTAGCATCCACC | | This study |
| Lys2972_ CaBM_Fw | TAACGGTAATGCAATTCCAGCAGGC | | This study |
| Lys2972_ CaBM _Fw | GCATCCACCCATGCAACC | | This study |
| Lys2972_ΔCBD_Fw | CACCACCACCACCACCAC | | This study |
| Lys2972_ΔCBD_Rv | GGCAGTATCAGCATAACGC | | This study |
| 2972_CaCl2_Fw | AGCATGGCTAACGGTGGAC | | This study |
| 2972_CaCl2_Rv | TGTTTTCACCTCGTTTTTCTTACC | | This study |
| CaBM_D219A_Fw | TCGTCGGCAGCGTCAGATGTGTATAAGAGACAGGGGATCCCAGTAGACATG | | This study |
| CaBM_D219A_Rv | GTCTCGTGGGCTCGGAGATGTGTATAAGAGACAGAGCCAAACCGTACCAAAC | | This study |
| pNZins_F | CGCTAAAACGTCTCAGAAAC | ^4^ | |
| pNZins_R | GTGATGGTTATCATGCAGGATTG | ^4^ | |
| T7 | TAATACGACTCACTATAGGG | Novagen | |
| T7 terminator | GCTAGTTATTGCTCAGCGG | Novagen | |
| ^a^ Abbreviations: Cam^r^, chloramphenicol resistance; Kan^r^, kanamycin resistance ^b^ | | | |

**Supplementary Table 1 references:**

1	Barrangou, R. *et al.* CRISPR provides acquired resistance against viruses in prokaryotes. *Science (New York, N.Y.)* **315**, 1709-1712, doi:doi:10.1126/science.1138140 (2007).

2	Lévesque, C. *et al.* Genomic organization and molecular analysis of virulent bacteriophage 2972 infecting an exopolysaccharide-producing *Streptococcus thermophilus* strain. *Applied and environmental microbiology* **71**, 4057-4068, doi:10.1128/aem.71.7.4057-4068.2005 (2005).

3	Hynes, A. P. *et al.* An anti-CRISPR from a virulent streptococcal phage inhibits *Streptococcus pyogenes* Cas9. *Nat Microbiol* **2**, 1374-1380, doi:10.1038/s41564-017-0004-7 (2017).

4	Hynes, A. P., Labrie, S. J. & Moineau, S. Programming native CRISPR arrays for the generation of targeted immunity. *mBio* **7**, doi:10.1128/mBio.00202-16 (2016).
